# Structure of biomolecular condensates from dissipative particle dynamics simulations

**DOI:** 10.1101/2019.12.11.873133

**Authors:** Julian C. Shillcock, Maelick Brochut, Etienne Chénais, John H. Ipsen

## Abstract

Phase separation of immiscible fluids is a common phenomenon in polymer chemistry, and is recognized as an important mechanism by which cells compartmentalize their biochemical reactions. Biomolecular condensates are condensed fluid droplets in cells that form by liquid-liquid phase separation of intrinsically-disordered proteins. They have a wide range of functions and are associated with chronic neurodegenerative diseases in which they become pathologically rigid. Intrinsically-disordered proteins are conformationally flexible and possess multiple, distributed binding sites for each other or for RNA. However, it remains unclear how their material properties depend on the molecular structure of the proteins. Here we use coarse-grained simulations to explore the phase behavior and structure of a model biomolecular condensate composed of semi-flexible polymers with attractive end-caps in a good solvent. Although highly simplified, the model contains the minimal molecular features that are sufficient to observe liquid-liquid phase separation of soluble polymers. The polymers condense into a porous, three-dimensional network in which their end-caps reversibly bind at junctions. The spatial separation of connected junctions scales with the polymer backbone length as a self-avoiding random walk over a wide range of concentration with a weak affinity-dependent prefactor. By contrast, the average number of polymers that meet at the junctions depends strongly on the end-cap affinity but only weakly on the polymer length. The regularity and porosity of the condensed network suggests a mechanism for cells to regulate biomolecular condensates. Interaction sites along a protein may be turned on or off to modulate the condensate’s porosity and tune the diffusion and interaction of additional proteins.

## I. INTRODUCTION

Biomolecular condensates are compositionally-diverse assemblies of protein in the cellular cytoplasm and nucleoplasm that form by liquid-liquid phase separation (LLPS).^1^ They have multiple roles in the cell^2^ including regulation,^3^ sequestering RNA stalled in translation during stress,^4^ cellular signaling,^5^ and organizing neuronal synapses in the brain.^6,7^ They also exert mechanical forces on chromatin suggesting that their material properties have biological significance.^8^ Although they lack a protective membrane barrier, their assembly and composition are tightly regulated by the cell, and loss of control is associated with ageing^9^ and chronic diseases including ALS, Alzheimer’s and Parkinson’s disease.^10^ They have been very well studied experimentally^11^ and are approaching the stage of being rationally tuned for industrial and therapeutic benefit.^12^ In spite of their importance in cellular physiology and disease, however, it is not yet understood how their material properties emerge from the molecular interactions of their constituents.

The main components of biomolecular condensates are intrinsically-disordered proteins (IDP) that have a little or no secondary structure,^13^ exhibit a large ensemble of conformations,^14^ and have little conserved sequence similarity.^15^ Instead of folding into a minimum-energy structure, IDPs sample many almost equi-energy conformational states in aqueous solution.^16,17^ As many of these states are extended, their conformational flexibility allows them to bind to multiple proteins via distributed, multivalent, reversible binding sites even at low concentrations.^18^ Their ability to explore interaction volumes whose size is comparable to the linear extent of the protein allows IDPs to phase separate from their surroundings into biomolecular condensates.

A wide range of experimental techniques have revealed the structure of BCs to resemble a fluid, mesh-like network in which the constituent IDPs transiently bind to each other but are locally disordered.^19^ Cryo-electron microscopy has shown that the condensed phase of FUS, an IDP associated with ALS, is amorphous, and lack internal ordered structures.^4^ Solution NMR experiments of Burke *et al*.^20^ show that FUS in its condensed phase is locally dynamic, making transient intermolecular contacts that are sufficiently short-lived that the proteins exhibit little or no secondary structure and remain disordered. At the same time, their fluorescence microscopy experiments show that the translational diffusion of FUS is much slower in the condensed phase than in the cytoplasm. A similar picture emerges for other IDPs. Ultrafast scanning Fluorescence Correlation Spectroscopy (FCS) shows that LAF-1 proteins inside P granules in C Elegans embryos form a surprisingly dilute phase that is two orders of magnitude below the dense phase concentration of the folded proteins lysozyme and γ-crystallin.^21^ The condensed network has a characteristic mesh size of 3 – 8 nm. Cryo-Electron Tomography of Sup35, a yeast prion protein, shows that it too forms a porous network with a mesh spacing around 10 nm.^22^ Non-biological peptides also phase separate into condensed droplets with an unexpectedly low internal density as a function of temperature and an inert crowding agent.^23^

Although *in vivo* BCs may contain hundreds of distinct protein components, as few as two,^6^ or even single protein species,^4^ are found to spontaneously condense *in vitro*. Banani *et al*.^24^ introduced a useful distinction in considering the role of the many components of BCs. They distinguished between a small number of IDPs - so-called *scaffolds* - that may be responsible for *forming* a condensed phase, and others - *clients* - that enter an existing phase to carry out biochemical functions. This has motivated research into identifying which molecular properties of the scaffold IDPs are required for them to undergo LLPS. It is perhaps surprising that IDPs should spontaneously condense into a low-density phase at all when it would appear *a priori* more favorable for them to remain dispersed and maximize their translational and conformational entropy. Researchers have drawn on ideas from polymer chemistry^16,25^ and applied a range of simulation techniques^26^ to understand how the molecular features of IDPs drive their phase behavior.

The material properties of biomolecular condensates evolve on length and time scales far above the atomic scale, and can exceed hours in *in vitro* experiments.^4^ This has limited atomistic Molecular Dynamics simulations to the conformational fluctuations of single molecules such as peptides,^27^ the disease-related proteins Huntingtin,^28^ α-synuclein,^29^ tau,^30^ and small oligomers.^31^ In an effort to go beyond this limitation, coarse-grained simulation techniques have been used in which the IDPs are simplified to linear chains of *beads* connected by *springs*.^32–34^ Reversible binding sites are represented as a direct attraction between selected beads or as an effective hydrophobic repulsion from the solvent. When the attractive beads are placed at the ends of the molecules they form so-called *telechelic* polymers.^35^ Brownian dynamics simulations of telechelic polymers show that they behave differently from pure random coils in dilute solution in that their radius of gyration decreases when their end-caps are attractive.^36^ Spatial correlations in the polymers’ conformations arise from the end-cap interactions, and the viscosity of the telechelic polymer solution is higher than that of a non-associating polymer solution. Telechelic polymers at the melt densities self-assemble into spherical or worm-like micelles in coarse-grained Molecular dynamics simulations.^37^ However, the high density of the melt phase is not representative of biomolecular condensates, which are typically found at low volume fraction *in vivo* and *in vitro*.^21^ Telechelics with hydrophobic end-caps self-assemble into micellar structures when their end-caps are strongly repelled from the solvent.^35^ Chatteraj *et al*. have used Langevin dynamics to explore the effects of polymer backbone flexibility and steric effects on the clustering of a model of membrane-bound nephrin, adaptor protein Nck1, and the actin-nucleating protein NWASP driven by interaction domains on the proteins.^34^ The system revealed a key relation between the polymers’ conformational flexibility and their ability to cluster: steric repulsion of inert sites located at the *termini* of the molecules reduced the mean cluster size more than inert sites near binding sites in the *middle* of the molecules. This effect was attributed to the terminal domains of the fluctuating polymers limiting access to the interior binding sites.

A fruitful picture of an IDP as a linear polymer with distributed binding sites – *stickers* – separated by flexible chain regions – *linkers* – was introduced by Harmon *et al*. in an illuminating study^38^ that used coarse-grained simulations to relate the solvation properties of disordered linker regions in model IDPs to their aggregation. On increasing their concentration, the IDPs were found either to pass through a phase transition and form a dense droplet, within which the proteins form a connected gel, or to directly pass from dispersed to the gel phase. Whereas stickers always act to promote aggregation, the effect of the linker regions depends on its interactions with the solvent. Hydrophobic linkers pull the stickers closer together, effectively increasing the inter-protein attraction, and so increase the propensity for phase separation. Hydrophilic linkers tend to swell the IDPs by attracting the solvent and push the binding sites farther apart, which increases the concentration needed for the gel phase to form. In subsequent work, they predicted that the differential solvation energy of the linkers in a mixture of four polymer types leads to structured droplets in which polymers more strongly repelled from the solvent are enclosed by those that prefer solvation.^33^ Molecular dynamics simulations have recently revealed a rich, sequence-dependent phase behavior in an amphiphilic polymer model of IDPs. The morphology and density of the condensed phase was elucidated as the hydrophobic fraction (*f_T_*) and its distribution along the polymer were varied for values *f_T_* > 0.6, which was the smallest value considered.^39^ The observed morphologies included spherical micelles, planar membranes, wormlike micelles, and a porous liquid phase.

Here, we study the equilibrium structure of a condensed phase of hydrophilic, polymers with self-associating *sticky* end-caps using Dissipative Particle Dynamics (DPD) simulations.^40,41^ This is a highly simplified model that reduces the molecular complexity of a biological IDP to a semi-flexible polymer with binding sites at its terminii. This apparently drastic reduction of complexity, however, retains a richness of behavior that appears to recapitulate much of the phase behavior of the original proteins. It provides a minimal model for the phase separation of IDPs that do not have a strong hydrophobic repulsion from the aqueous solvent. The model IDPs studied here differ significantly from those in two recent simulation studies. Firstly, in the nomenclature of Harmon *et al*.,^38^ who studied the effects of the solvation of linker regions on the IDP’s phase separation, the backbone of the polymers in our study is always highly solvated. Secondly, the polymers possess no hydrophobic regions in contrast to the work of Statt *et al*.^39^ who studied the aggregation of IDPs containing a minimum fraction *f_T_* ≥ 0.6 of hydrophobic beads. We find that polymers with weak endcap affinity form box-spanning networks as observed previously.^38^ But, surprisingly, polymers with sufficiently *sticky* end-caps phase separate at very low concentrations into porous droplets resembling three-dimensional networks. The end-caps reversibly bind at junctions whose spatial separation is *independent* of the polymer concentration, and insensitive to their end-cap affinity over a wide range (above a threshold value for which a stable network forms). The polymers diffuse slowly within the networks and exchange with those dispersed in the solvent. Their formation is reminiscent of the two-dimensional phase separation that produces domains in lipid membranes and provides a conceptual foundation for understanding biological membranes.^42^

DPD is a coarse-grained, explicit-solvent technique^40,41^ that has been widely used to simulate equilibrium properties of soft matter systems. These include the phase morphologies of polymers,^43^ polymer microphase separation,^44^ telechelic polymers,^35^ and lipid bilayer membranes.^45,46^ It has also been used to explore non-equilibrium phenomena such as domain formation in vesicles,^47^ the fusion of vesicles and membranes,^48^ the self-assembly of amphiphilic vesicles,^49^ and the interaction of nanoparticles with membranes.^50^ An extensive comparison of the equilibrium properties of amphiphilic membranes on length scales much larger than the constituent molecules showed that they are consistent for different coarse-graining strategies,^51^ a result that gives confidence in applying such techniques to polymeric aggregates. The reader is referred to a recent perspective article for the latest methodological developments of DPD.^52^

Our results reveal a surprisingly rich phase behavior for hydrophilic polymers with sticky end-caps as their length and end-cap affinity are varied. It is likely that this richness contributes to the biological functions of biomolecular condensates. We hypothesize that the independence of the network’s porosity (as measured by the inter-junction separation) on the polymers’ binding affinity and concentration, and the scaling of the inter-junction separation with the end-cap separation may be used by cells to control biochemical activity inside biomolecular condensates. Selectively activating binding sites on the scaffold IDPs, via diffusing kinases for example, will modulate the porosity of their condensed phase and influence the recruitment and interactions of additional proteins.

The paper is organized as follows: in Section II we summarize the DPD simulation method and the polymer model we use to represent IDPs. Section III presents our results and quantifies the internal structure of the condensed phase. Finally, Section IV contains our conclusions and identifies potential biological and therapeutic applications of the observed relations between the structure of the condensed phase and its constituent molecular properties.

## II. SIMULATION METHOD

In this work, we study the phase behavior and structure of the condensed phase of telechelic polymers (Fig. 1) as a simplified model of biomolecular condensates. In order to reach the length and time scales on which the condensed phase equilibrates, we use the coarse-grained molecular simulation technique of Dissipative Particle Dynamics (DPD).^40,41^ The *beads* in DPD represent small volumes of atoms or molecular groups and interact via three short-ranged, soft, momentum-conserving, pairwise-additive forces. Once the forces are defined, Newton’s laws of motion are integrated to generate bead trajectories over time. The advantage of DPD over conventional Molecular dynamics simulations for simulating complex fluids is that the softer potentials permit a large time-step and very long simulation times. The momentum-conserving forces also retain the correct hydrodynamic behavior of a fluid, which distinguishes it from Brownian dynamics in which all motion is diffusive.

**Figure 1.**
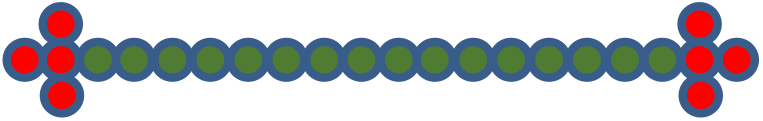
Cartoon of the polymer E_4_ B_16_ E_4_ that represents a generic IDP in the simulations. We do not attempt to map a particular protein’s residue sequence in the model, nor include sequence-dependent secondary structure. The IDP is reduced to a semi-flexible polymer composed of hydrophilic backbone beads B with two hydrophilic, self-associating end-caps composed of beads E. The backbone length is varied to represent proteins of different molecular weight and the end-cap interaction determines their binding affinity. The end-cap shape is chosen to create an approximately isotropic interaction volume.

The conservative force between beads i and j takes the form:

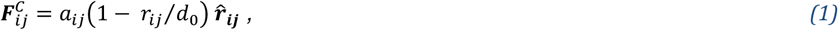

for *r_ij_* < *d*_0_. The maximum value of the force is *a_ij_*; ***r_ij_*** = ***r_i_ – r_j_*** is the relative position vector from bead j to bead i, *r_ij_* is its magnitude, and 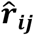 is the unit vector directed from bead j to bead i. The conservative force parameter *a_ij_* gives beads a chemical identity, such as oily hydrocarbon chains that are hydrophobically repelled from water.

The other two non-bonded forces constitute a thermostat that ensures the equilibrium states of the simulation are Boltzmann distributed.^41^ The dissipative force is:

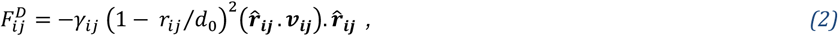

where *γ_ij_* is the strength of the dissipative force and ***v_ij_*** is the relative velocity between beads i and j. This force destroys relative momentum between interacting particles. The random force is:

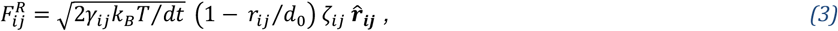

where k_B_T is the system temperature. This force creates relative momentum between pairs of interacting particles. Here, *ζ_ij_* is a symmetric, uniform, unit random variable that is sampled for each pair of interacting beads and satisfies *ζ_ij_* = *ζ_ij_*, 〈*¶_ij_*(*t*)〉 = 0 and 〈*ζ_ij_*(*t*)*ζ_kl_*(*t′*)〉 = (*δ_kl_δ_jl_* + *δ_il_δ_jk_*)*δ*(*t* – *t′*). The pairwise forms of the dissipative and random forces conserve momentum. The factor 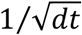 is required in the random force so that the discretized form of the Langevin equation is well defined.^41^

Beads are connected into polymers using Hookean springs with potential energy:

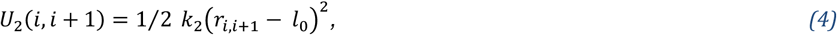

where the spring constant, *k*_2_, and unstretched length, *l*_0_ may be different for each bead type pair i, j. The semi-flexible nature of polymers is represented by a chain bending potential applied to the angle θ defined by adjacent bead triples:

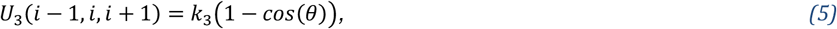

where the stiffness parameter, k_3_, may be different for different bead triples. Table 1 lists the values of all the non-bonded and bonded parameters.

**Table 1.**
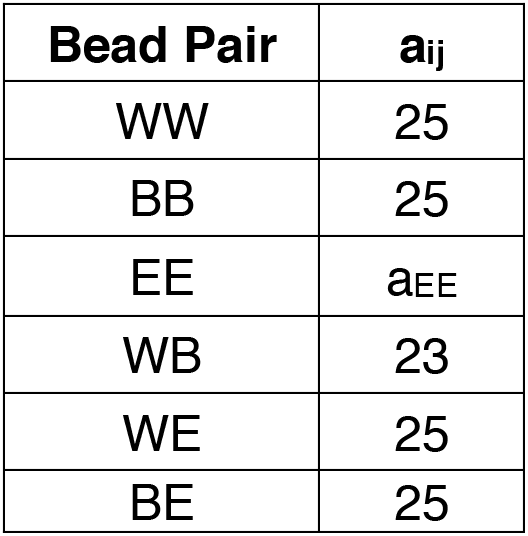
Non-bonded conservative interaction parameters a_ij_ for all bead types (in units of *k_B_T*/*d*_0_). The backbone (B) and endcap (E) beads are hydrophilic, which represents a polymer in a good solvent, and the parameter a_EE_ is varied to modify the end-caps’ binding affinity: smaller values of aEE correspond to increased attraction between the E beads as described in the Supplementary Material section 1. The reduced value of a_WB_ ensures that the polymer backbone remains solvated in the network phase. The dissipative force parameters are 4.5 for all bead pairs (in units of 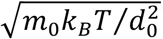). Beads are connected into polymers using Hookean bonds. Based on previous simulations of amphiphilic membranes,^45^ the bond potential constrains the bonds’ mean length, and the same values, 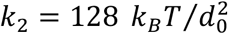 and *l*_0_ = *d*_0_/2, are used for all bonded beads (EE, EB, BB). Chain stiffness is imposed by a bending potential for all BBB triples along the backbone with bending constant *k*_3_ = 5 *k_B_T*.

Simulations are performed in the NVT ensemble in which the number of particles, system volume and temperature are constant. Most simulations take place in a cubic box with linear size *L* = 48*d*_0_, where *d*_0_ is the range of the non-bonded DPD forces and defines the length-scale in the simulations. Periodic boundary conditions are applied in all three dimensions. The bead density 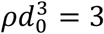, so the total number of beads is *ρL*^3^ = 331,776. A larger simulation box *L* = 64*d*_0_ is used as noted in the text to check the system size dependence (see Supplementary Material). All beads have the same mass, m_0_, and simulations take place at reduced temperature *k_B_T* = 1, so that simulation time is measured in units of the DPD time-scale 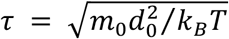. The integration time-step is chosen to be 0.02 *τ*. A fixed number of IDPs are initially distributed randomly throughout the simulation box together with sufficient water beads to give the prescribed density, and the simulation is allowed to evolve to equilibrium after which observables are sampled for analysis (see Section 4 of the Supplementary Material for the error analysis). The evolution of the networks is very slow, particularly for longer polymers. We discard at least 3,000,000 time steps before ensemble averages are constructed by sampling over a further 1,800,000 time steps. A relevant time-scale can be assigned to the simulation by comparing the dimensionless diffusion constant of polymers in the equilibrium droplet, 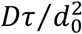, with experimental values. Burke *et al*. report the diffusion constant of the disordered FUS N-terminal domain within its phase separated state to be 0.4 μm^2^/sec.^20^ We take the DPD bead size to be *d*_0_ = 1 *nm*, in line with previous work on amphiphilic membranes,^45^ so that one DPD bead represents approximately 3 amino acid residues. The DPD time-scale is then *τ* = 9 *ns*, and a typical run of four million steps represents 720 microseconds of real time. The precise value depends on the polymer molecular weight, but as we are interested in equilibrium properties, we do not attempt to fix the time-scale more precisely.

## III. RESULTS AND DISCUSSION

### A Telechelic polymers spontaneously aggregate at low concentrations

We study the formation and structure of the condensed phase of a model IDP by representing it as a linear polymer with *sticky* end-caps (Fig. 1). The molecular architecture is described by the formula E_M_B_N_E_M_ where N is the number of backbone beads of type B, which is varied to represent IDPs of different molecular weight, and each end-cap contains M beads of type E. Each backbone bead (B) corresponds to several amino acid residues, and the end-cap E beads represent nonspecific, reversible interaction domains. The aqueous solvent is represented by a single bead W. We note here that the backbone beads do not distinguish between different types of amino acid and both backbone and endcap beads are hydrophilic so there is no hydrophobic repulsion from the solvent driving the aggregation. The attraction between the end-caps is quantified by a dimensionless binding affinity *ϵ* that is defined so that *ϵ* = 0 corresponds to no affinity and *ϵ* = 1 is a strong affinity. The definition of the binding affinity in terms of the DPD conservative force parameters and further details of the simulations and are given in Section 1 of the Supplementary Material.

Preliminary simulations showed that IDPs whose end-caps have M < 4 do not assemble into a condensed phase even for strongly attractive end-caps (data not shown). All IDPs therefore have end-caps with M=4 as shown in Fig. 1, and are described by the formula E_4_B_N_E_4_. The cross shape of the four end-cap beads is chosen so that they expose an approximately isotropic interaction surface. Hereafter, we refer to polymers E_4_B_N_E_4_ with N backbone beads by the formula B_N_ for brevity. Early simulations also showed that large conformational fluctuations of completely flexible polymers concealed their binding sites within compact conformations and precluded aggregation, an effect reported previously in the literature.^34^ Therefore, a bending stiffness is applied to the polymer backbone to represent the semi-flexible nature of IDPs without assuming any particular secondary structure (see Table 1).

We first describe the conditions under which polymers of type B_16_ spontaneously aggregate into networks as their concentration, backbone length and end-cap affinity are varied. The polymer concentration is defined as the ratio of the number of polymers to the total number of molecules (polymer + solvent), and is therefore the number fraction. Figure 2 shows snapshots from simulations of 634 polymers of type B_16_ in a simulation box of size (48d_0_)^3^ with the affinity increasing from left to right and top to bottom. Unless otherwise stated, this system size is used for all the results presented. Within the condensed phase, the polymer end-caps meet at *junctions* while a small number of polymers remain free in the solvent.

**Figure 2.**
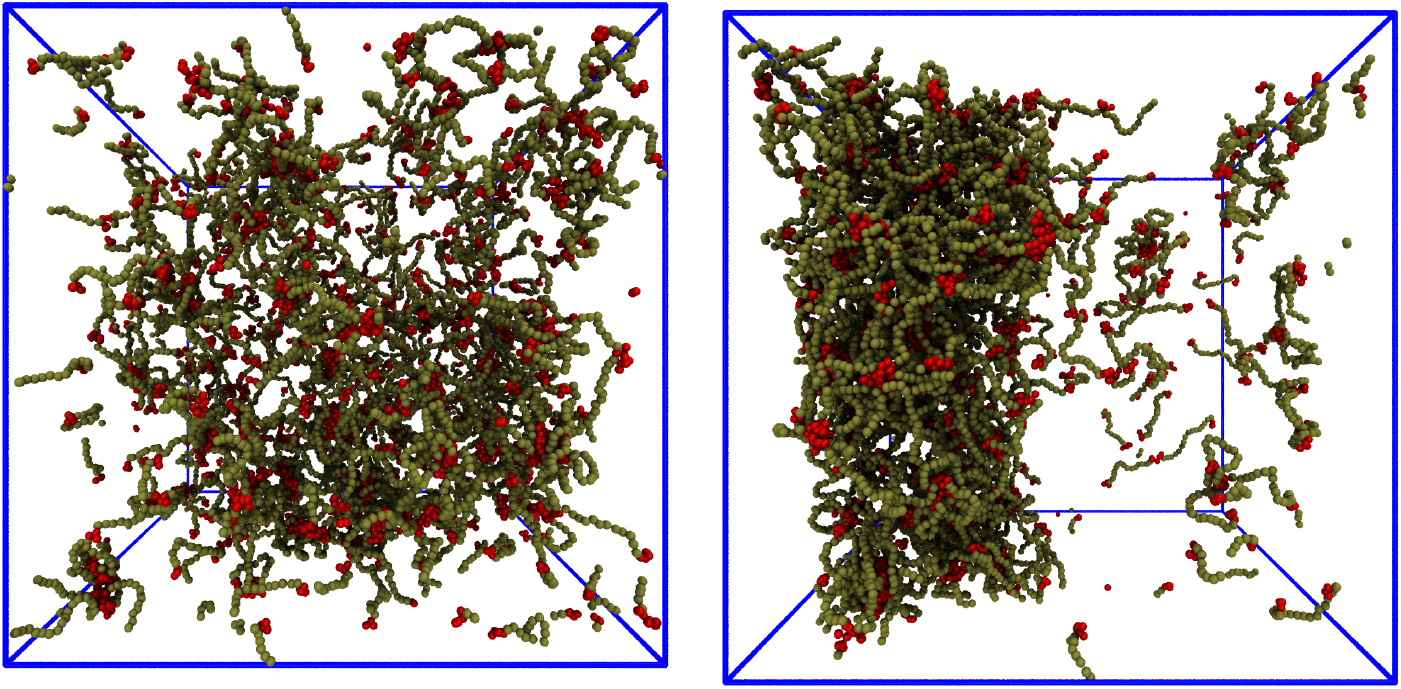
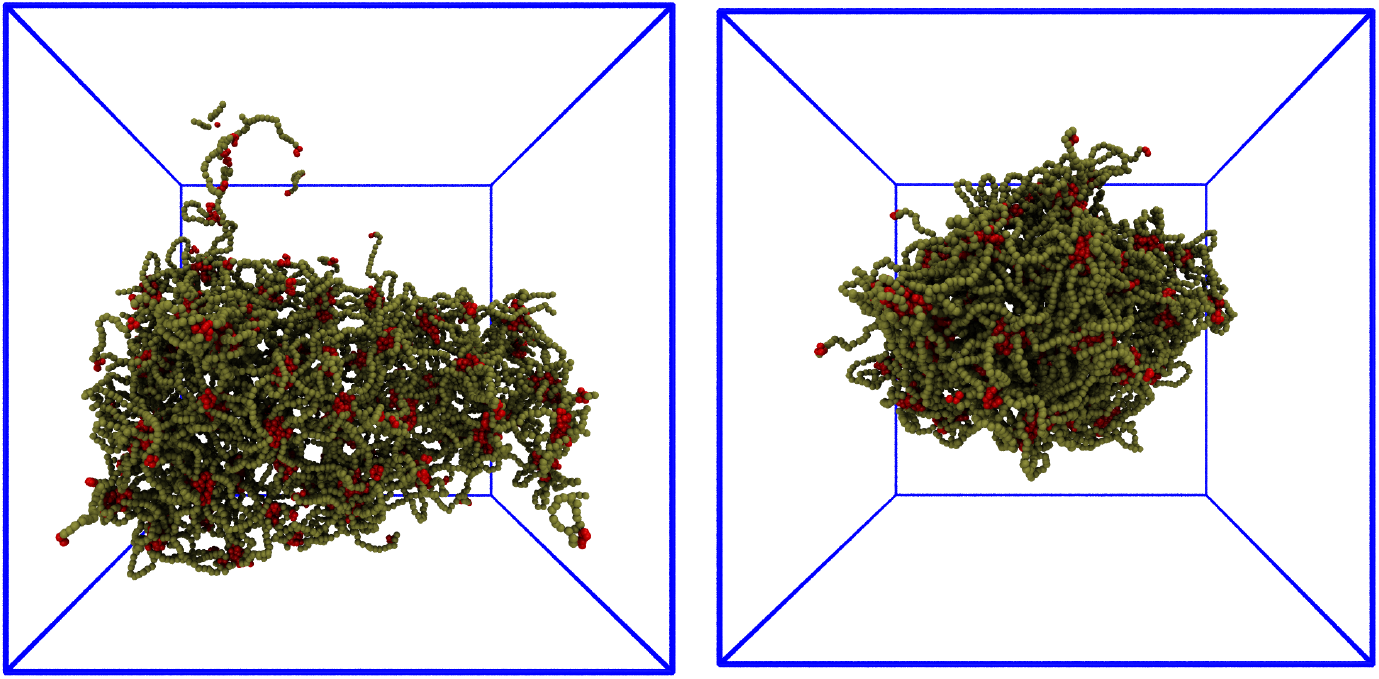
Equilibrium configurations of E_4_B_16_E_4_ polymers in water (invisible for clarity in all snapshots) at a concentration of 0.002 (634 polymers) for four affinities spanning the range from a dispersed phase to a condensed network. Polymers with a weak end-cap affinity (ε = 0.72, top left) form transient clusters in equilibrium with dispersed polymers and a percolating network at higher concentrations without phase separation. Polymers with stronger affinities, (ε = 0.76, top right, and ε = 0.8, 0.84 bottom left and right) phase separate at low concentrations. The apparently disconnected polymers are a consequence of the periodic boundary conditions.

Figure 2 shows that the junctions (colored red in the figure) appear to have a somewhat regular distribution in space, but do not form a lattice structure. We quantify this observation in the next section. Similar behavior is found for polymers of all lengths simulated - B_8_, B_10_, B_24_, B_32_ (see Fig. S1 and supplementary movies 4 and 5). No polymers in this length range with an endcap affinity below ε = 0.6 were found to form a network. Henceforth, we refer to affinities close to ε ~ 0.6 as *low*, values at or above ε ~ 0.8 as *strong*, and values above ε ~ 0.88 as *very strong* with the understanding that these are qualitative designations only. They reflect the observed differences in the appearance of networks seen, for example, in Figures S1 - S3.

Visual inspection of the simulation snapshots shows that polymers continually join and leave a network, and it can break up and reform during a simulation, especially for lower affinities. This raises the question of which network to use for quantitative measurements. Condensed phases of IDPs in experiments are often microns in size, while our largest simulated networks approach 50 nm. In the remainder of this work, we report quantitative properties only for the single largest network in a simulation as it evolves in time. The structural properties of simulated networks containing hundreds to thousands of polymers are independent of their size indicating that they form a thermodynamic phase. This is referred to as the *Largest Equilibrium Network* (LEN). The algorithm for identifying the LEN is described in Section 2 of the Supplementary Material.

Figure 3 shows that the number of polymers in the LEN increases slightly sub-linearly with concentration above an end-cap affinity-dependent threshold. Polymers with weaker affinities (Figure 2, top left, ε = 0.72 and supplementary ovie 1) remain dispersed in the solvent until their concentration is sufficiently high for them to form a network occupying the whole simulation box. Polymers with higher affinities (Figure 2, top right, ε = 0.76, bottom left, ε = 0.8, and supplementary movie 2; Figure 2, bottom right, ε = 0.84, and supplementary movie 3) phase separate at very low concentration into a porous network. These distinct structures arise from the balance of the enthalpy gain of polymers binding to each other and the entropy of translational and conformational fluctuations of the dispersed polymers. Polymers with strong end-cap affinity phase separate because their conformational entropy is *largely unchanged* in the condensed phase and the enthalpy gain of binding lowers the system’s total free energy (see Section C). For example, the LEN composed of B_16_ polymers with high-affinity end-caps (e.g., ε = 0.8) in a system with a concentration 0.002 contains almost all of the available polymers (622 polymers out of 634). Polymers with lower affinities (e.g., ε = 0.68) remain dispersed until their concentration is sufficiently high for them to form a box-spanning network because the (weaker) enthalpy gain of their end-caps binding at junctions is insufficient to overcome their loss of translational entropy.

**Figure 3.**
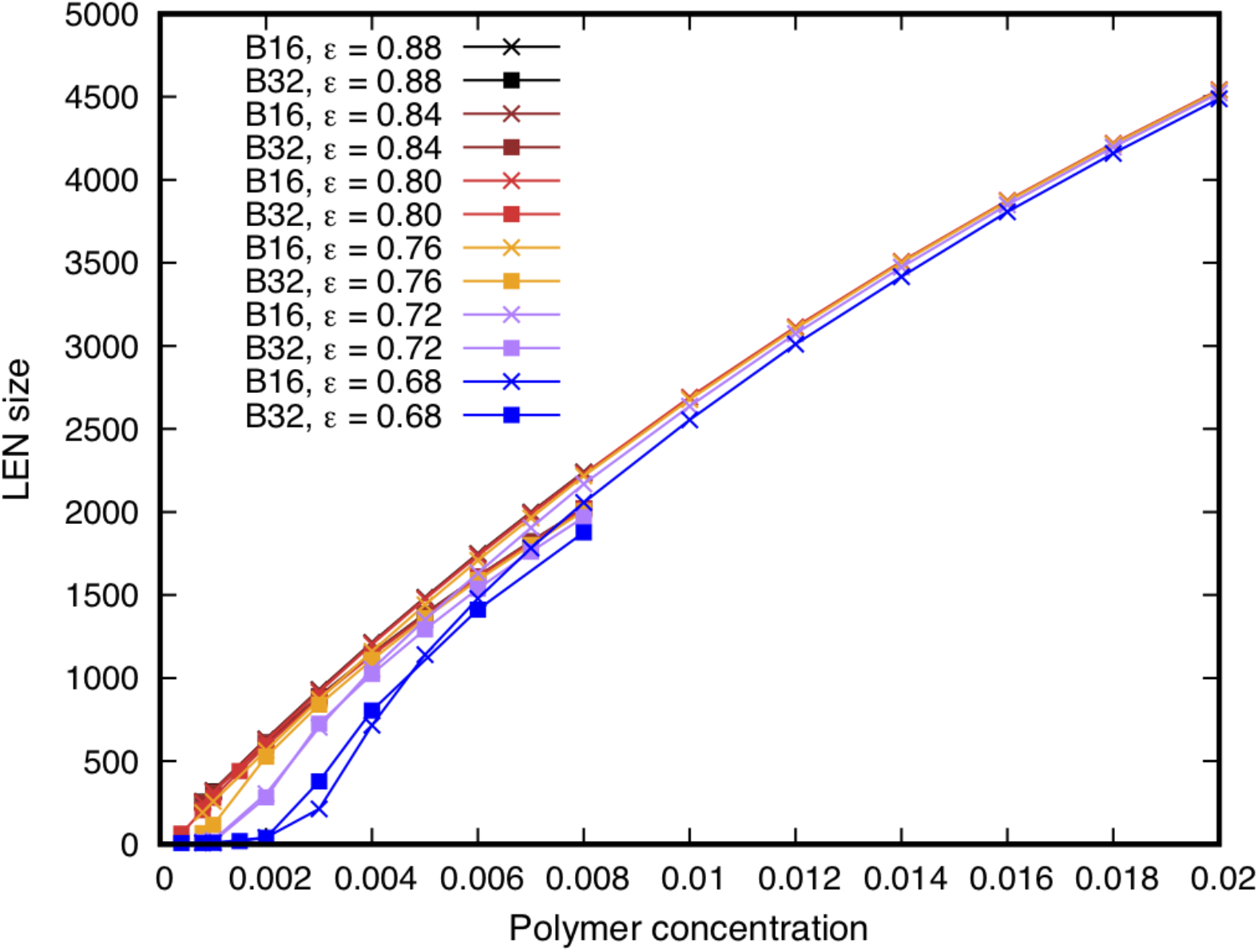
The number of E4BNE4 polymers in the Largest Equilibrium Network as a function of the polymer concentration for backbone lengths N = 16, 32 and end-cap affinities ranging from weak (lower curves) to strong (upper curves). The network’s size increases linearly with polymer concentration for all affinities above a threshold (ε ~ 0.76), and becomes largely independent of the polymer length and end-cap affinity at higher concentrations. The qualitative distinction between phase separation of high-affinity polymers and formation of a percolating network for low affinity polymers at high concentrations seen in Figure 2 is reflected here as polymers with weak end-cap affinity (ε = 0.68) are unable to aggregate at low concentrations because the enthalpic gain of binding is insufficient to overcome their translational and conformational entropy. Note that the curves for higher affinities overlap closely. Statistical errors are of the same order as the symbol size.

We reiterate here that the simulated polymers are hydrophilic and their backbones do not strongly interact with each other. This can be seen in Figure 2 where the phase separated network is visibly porous. Harmon *et al*.^38^ have observed a similar distinction between phase separation or reversible gelation without phase separation in lattice Monte Carlo simulations of fixed-length polymers composed of attractive, sticker regions separated by flexible linkers. In their work, the solvation properties of the linker regions between attractive sites were varied between highly solvated and less solvated. They observed that highly solvated, and hence swollen, linkers drive the polymers to form a reversible gel at sufficiently high concentrations. However, linkers with a small solvation volume promote binding of the attractive sites by bringing them closer together, leading to phase separation before gelation. Intriguingly, however, phase separation appears in our system in what corresponds to the *large solvation volume* limit in their model, for which they observed reversible gelation. We discuss this phase transition further in Section D. The observed phase separation appears for different combinations of polymer length and end-cap affinity, and it is natural to ask how the internal structure of the condensed phase depends on the polymer architecture. We turn to this question in the next section.

### B Junction separation in the network scales with the polymer length independently of the polymer concentration and end-cap binding affinity

Figure 2 and Figure S1 show that the junctions appear rather regularly distributed within the condensed networks. We quantify this observation by defining the distance between connected junctions as the mean value of the end-to-end length of all polymers whose end-caps reside on the junctions. This quantity is averaged over all pairwise-connected junctions in the LEN to give the *Mean Junction Separation* L_ee_. Close visual inspection of the networks reveals that some polymers have both of their end-caps residing on the same junction (see Figs. S2, S3, S6 and supplementary movie 4). We exclude these ring-like conformations from the measurements of L_ee_ because they do not contribute to the junction separation (see Section 3 of the Supplementary Material for details). Polymers with very strong affinity adopt ring-like conformations in the dispersed phase even at very low concentrations (Fig. S2).

Figure 4 shows the variation of L_ee_ with polymer concentration for networks composed of polymers B_8_ to B_32_ whose end-caps have very strong affinity (ε = 0.96), strong affinity (ε = 0.8) and weak affinity (ε = 0.68). We note here that all quantitative results presented are sampled from equilibrium states of the network after discarding at least the first two million time-steps. We have verified that the network is fluid, which allows it to evolve towards an equilibrium state, by performing a simulated Fluorescence Recovery after Photobleaching Experiment (FRAP)^53^ (Fig. S5). A network of 1215 B_16_ polymers was allowed to form and equilibrate. The fluorescence bleaching effect in a FRAP experiment is simulated by assigning a different display color to the end-cap beads of all polymers in one half of the network (yellow beads in Fig. S5). The labelled polymers are subsequently observed to diffuse through the network. Networks of B_8_ and B_24_ polymers show similar behavior as do networks of polymers with lower affinity (data not shown). Further discussion of the statistical errors is given in the Supplementary Material (Figs. S7, S8). We confirm that L_ee_ is independent of the system size by performing some simulations in a larger box (64d_0_)^3^ (Figs. S9, S10).

**Figure 4.**
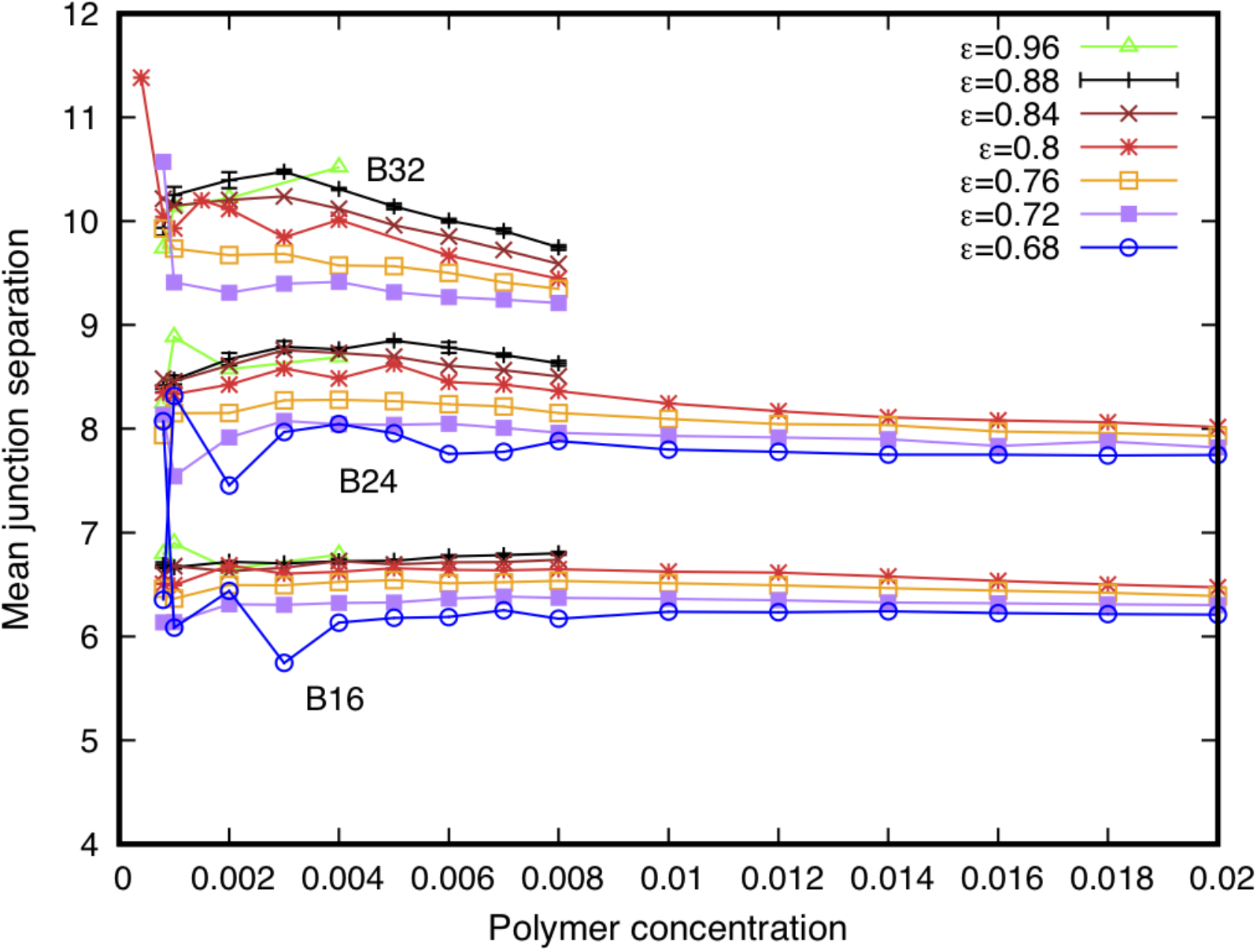
The major structural property of the Largest Equilibrium Network is the mean junction separation shown here as a function of polymer concentration for a range of backbone lengths and affinities. The junction separation in networks of polymers of all backbone lengths and end-cap affinities studied is independent of the polymer concentration above a minimum threshold concentration. The asymptotic value of the separation depends systematically on the backbone length, but only weakly on the end-cap affinity. The large fluctuations in the junction separation for polymers B_16_, B_24_, B_32_ at concentrations around 0.001 are due to the instability of the small networks. Statistical errors are of the same order as the symbol size and are only shown for ε = 0.8. Systematic errors are discussed in the Supplementary Material.

Three interesting observations can be made from this data. First, the mean junction separation Lee in the LEN is independent of polymer concentration above a minimum value at which a stable LEN is formed. This indicates that the internal environment of the condensed phase experienced by the polymers, as measured by Lee in Fig. 4, is structurally well-defined and similar despite the variety of morphological states in which they are situated ranging from nearly-spherical droplets to extended structures spanning the periodic boundaries of the simulation box (*cp*. Fig. 2).

Second, L_ee_ is almost independent of the end-cap affinity from the lowest value (ε = 0.68) to the highest value studied (ε = 0.96) as shown by the near superposition of the curves in Fig. 4 for polymers with the same backbone length but different affinities. As no networks were observed for polymers with affinities below ε = 0.68, it appears that as soon as the affinity/backbone length permit a stable network to form, the internal geometric structure of the network remains unchanged for higher affinities. Taken together, these results show that the supramolecular properties of the LEN are independent of its size and shape.

Third, some polymers adopt ring-like conformations in the LEN at all concentrations studied. The fraction of polymers in the LEN that form rings decreases with increasing polymer concentration to an asymptotic value that depends on the polymer length but only weakly on the end-cap affinity (Fig. S4). This value is about 10% for polymers with low affinity (ε = 0.68) end-caps and 15-20% for those with high affinity (ε = 0.8) end-caps. Shorter polymers more easily adopt ring-like conformations, and those with the highest affinity studied (ε = 0.96) have the largest fraction of ring conformations, above 50%, for *low* concentrations of polymers B_24_ and shorter (see top curves in Fig. S4). The fraction of rings in networks composed of the long polymers B_32_ and B_48_ with the highest affinity studied (ε = 0.96) falls rapidly to similar values for those of low-affinity polymers once their concentration exceeds 0.002, and less than 15% adopt ring conformations in the networks. The persistence of these rings in networks at higher concentrations suggests that they are not a significant factor in driving the dispersed polymers into the network phase. This distinguishes the networks from the theoretical transition between ring-like polymers and a network phase analyzed by Semenov, Nyrkova, and Cates.^54^

### C The separation of connected junctions shows self-avoiding walk scaling with the polymer length

The near superposition of the curves of L_ee_ in Figure 4 for polymers with the same backbone length N but different concentrations and affinities suggests that the mean junction separation L_ee_ may scale with the polymer backbone length. Figure 5 shows that indeed L_ee_ / N^ν^ is independent of polymer length for all affinities studied from weak (ε = 0.68) to very strong (ε = 0.96). The power v = 0.6 is the Flory exponent for self-avoiding random walks (SAW).^25^ This scaling relation holds for both phase-separated droplets and box-spanning networks that form without phase separation. The prefactor of the scaling relation has a weak affinity dependence as the curves move down the ordinate axis with decreasing affinity. The scaling is independent of time for both high (Fig. S7) and low (Fig. S8) end-cap affinities showing that it is an equilibrium property. This result indicates that polymers within the condensed network phase exhibit the same SAW conformational fluctuations as dispersed polymers. Note that only polymers with the strongest affinity studied (ε = 0.96) are observed to phase separate for the long B_48_ polymers. We include this case here to demonstrate the robustness of the scaling relation but do not discuss other quantitative measures for B48 systems as their slow dynamics makes reaching equilibrium computationally expensive.

**Figure 5.**
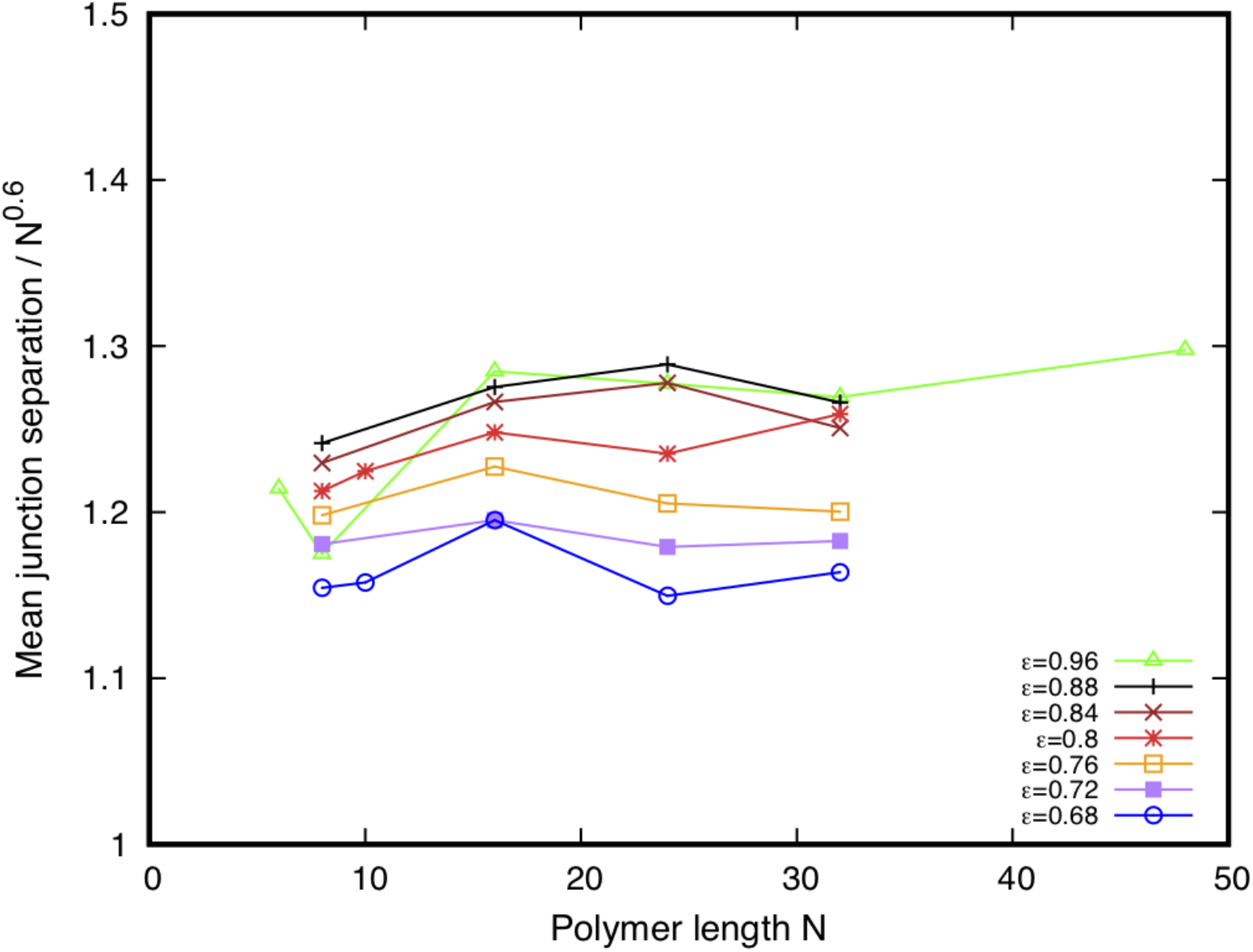
The mean junction separation in the Largest Equilibrium Network scales with the polymer backbone length as a self-avoiding random walk with Flory exponent 0.6 for all the end-cap affinities studied from weak (ε = 0.6), to strong (ε = 0.8), and very strong (ε = 0.96). A small affinity-dependent prefactor is evident by the displacement of the curves down the ordinate axis as the affinity decreases. This dependence of the junction separation on polymer length indicates that the networks are sufficiently porous that the polymer backbones fluctuate as self-avoiding polymers while their end-caps reversibly bind at the network junctions. Note that only the highest affinity (ε = 0.96) permits polymers of length B_48_ to phase separate.

### D Polymer distribution at the junctions depends on end-cap affinity

An important function of biomolecular condensates is to concentrate biochemical reactions.^2^ This requires that interacting proteins are able to diffuse within the condensate in order to react. The robust scaling of the junction separation with the polymer’s backbone length in the simulated droplets implies that their porosity depends only on the effective separation of the polymers’ binding sites and is independent of their concentration and affinity. In networks such as those shown in Figure 2, the polymers diffuse within the network and exchange between the network and the dispersed phase. Although the equilibrium properties of the network do not change in time, there is no thermodynamic requirement for them to be spatially uniform. The distribution of polymers among the junctions of the network influences its porosity which, in turn, determines how easily additional molecules are able to diffuse through the network and encounter each other to undergo biochemical reactions.

We refer to the number of polymers whose end-caps meet at a given junction as the *junction mass*. The B_16_ polymers phase separate into a condensed network at very low concentrations (< 0.002). Figure 6 shows the variation of the junction mass in a system of B_16_ polymers with increasing concentration for end-cap affinities from weak (ε = 0.6) to strong (ε = 0.88). The junction mass for weak-affinity end-caps (ε ≤ 0.72) monotonically increases with increasing polymer concentration but undergoes no apparent transition. By contrast, the junction mass for B_16_ polymers with stronger affinities (ε ≥ 0.76) exhibits a flat regime, which lasts until a concentration of around 0.01, before increasing for higher concentrations. The flat portions of the mean junction mass curves indicate the coexistence region in which the droplet is in equilibrium with the surrounding dilute phase similar to the flattened isotherms in the pressure/density indicator diagram of a fluid below its critical temperature. The width of the coexistence region decreases with decreasing affinity. When the droplet has grown to the size of the simulation box, the mean junction mass increases with a further increase of concentration.

**Figure 6.**
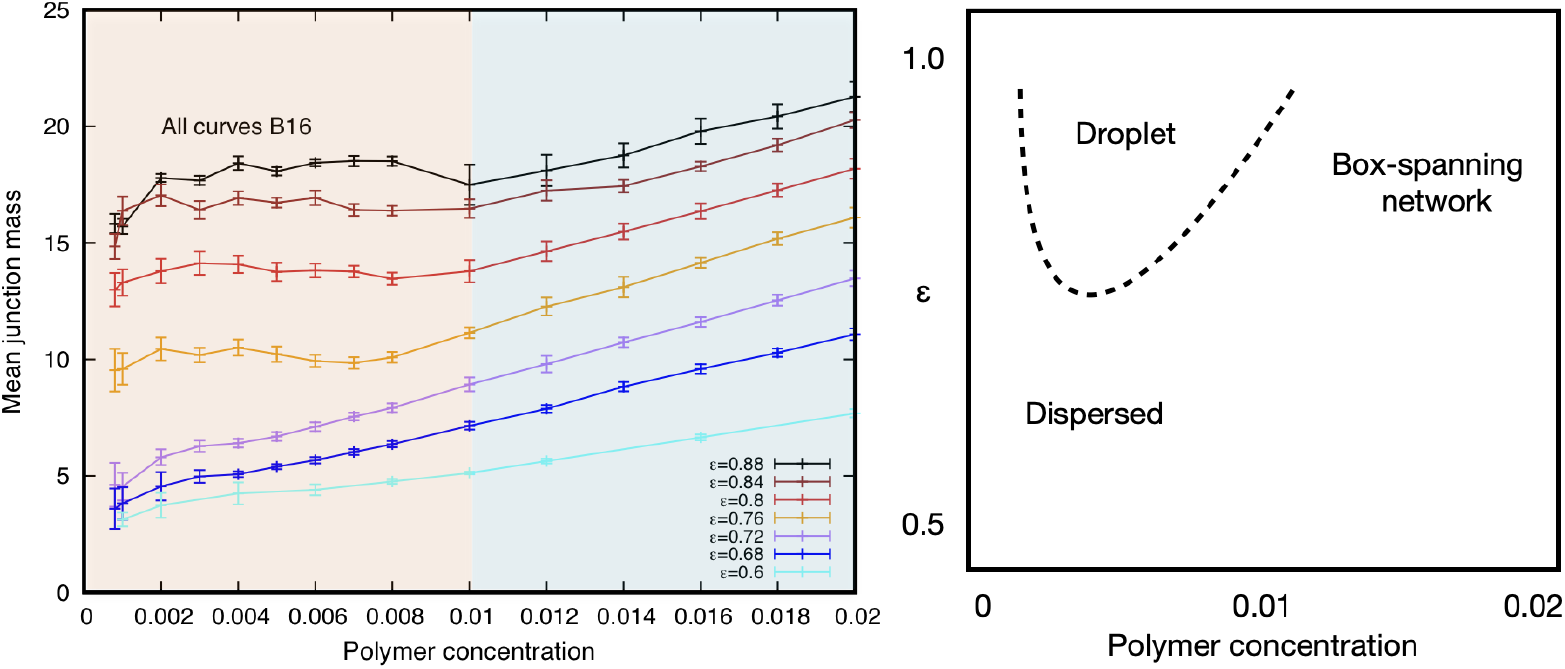
(Left) The mean number of B_16_ polymers meeting at the network junctions (the junction *mass*) for end-cap affinities from weak (ε = 0.6, bottom curve) to strong (ε = 0.88, top curve). The junction mass increases monotonically with increasing concentration for low affinities. Above a critical affinity (ε ≥ 0.76), the curves exhibit a flattened region before rising with concentration. The flat regions delimit the coexistence region (within the left-hand shaded region) in which polymers phase separate into a droplet from the single-phase region (right-hand shaded region) in which the junction mass increases with concentration. The statistical error bars are calculated from three consecutive runs of 600,000 steps each. (Right) Qualitative phase diagram for B_16_ polymers in the ε/concentration plane showing that the system phase separates at low concentrations into a dense droplet coexisting with a dilute phase for affinities above a critical affinity (ε ~ 0.76).

The right-hand panel in Figure 6 shows the qualitative phase diagram of B_16_ polymers in the ε/concentration plane. This is a cut through the threedimensional phase space in which the polymer length forms the third axis. At very low concentrations, the polymers are dispersed in the solvent and form a single phase. On increasing the concentration, polymers with affinities below the critical value remain in a single, dispersed phase until the transient clusters merge into a single box-spanning network. Polymers with affinities above the critical value undergo phase separation for which the low-concentration phase boundary depends only weakly on the end-cap affinity. As the concentration increases, the condensed phase contains an increasing fraction of the polymers until, at a polymer length-dependent upper concentration, the condensed phase forms a single, box-spanning phase that contains all the polymers. The low-concentration arm of the two-phase region is hard to locate precisely because polymers with affinities above the critical value tend to spontaneously aggregate even at very low concentrations (*cp*. Fig. S2).

Similar, although weaker, behavior is seen for longer polymers. Figure 7 shows the mean junction mass for systems of B_24_ and B_32_ polymers across the same affinity range as Figure 6. There is a slight flattening of the curves for B_24_ polymers with strong affinities (ε ≥ 0.8) while the curves for weaker affinities increase monotonically. The curves for B_32_ polymers show no obvious flattening for all affinities studied. The phase transition is pushed to higher end-cap affinities as the polymers increase in length. Comparing the width of the coexistence regions in Figures 6 and 7 shows that the droplet phase boundaries shrink with increasing polymer length. The low-concentration arm drops to very low values and the high-concentration arm also moves to lower values. This suggests that IDPs such as α-synuclein (140 residues) would spontaneously aggregate at extremely low concentrations if they only possessed binding sites at their terminii. In order to be soluble at physiological concentrations, yet still undergo phase separation at higher concentrations, they must have multiple, weak binding sites whose separation along the protein backbone is not too large. This arrangement of binding sites is found for IDPs.^18^

**Figure 7.**
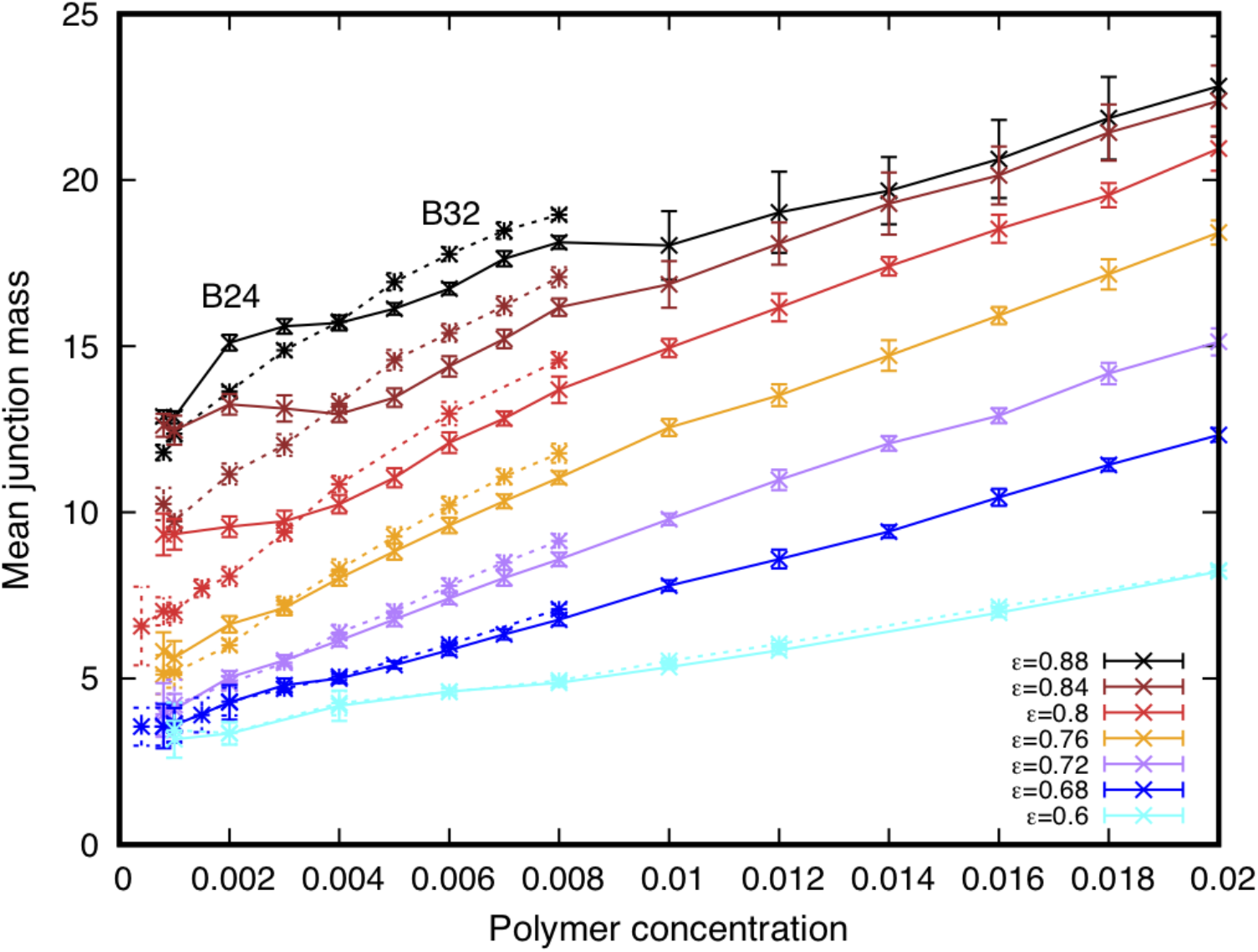
The junction mass for polymers B_24_ (solid) and B_32_ (dashed) with end-cap affinities from strong (ε = 0.88, top curve) to weak (ε = 0.6, bottom curve). For clarity, only the ε = 0.88 curves are labelled with the polymer length. A slight flattening of the B_24_ curves seen for concentrations between 0.002 and 0.004 for the strongest three affinities is an indication of a possible phase transition, but it is much weaker than that seen in Figure 6 for B_16_ polymers. This indicates that the longer polymers require a higher end-cap affinity to phase separate. Statistical error bars are obtained from three consecutive runs of 600,000 steps each.

The mean junction mass does not fully reflect the distribution of polymers among the network’s junctions, particularly if it is non-Gaussian. Figure 8A shows that the junction mass distribution for high-affinity networks (B_16_, ε = 0.8) is very broad, ranging from 3-30 polymers/junction, and approximately Gaussian. The two histograms are sampled from widely-separated times in the simulation and show that the mass distribution is invariant over time although polymers diffuse through the network (as confirmed by the FRAP simulation shown in Fig. S5). The enhancement in the number of junctions with fewer than 5 polymers arises because junctions near the surface of the network are more surrounded by solvent compared to those in the network’s core.

**Fig. 8.**
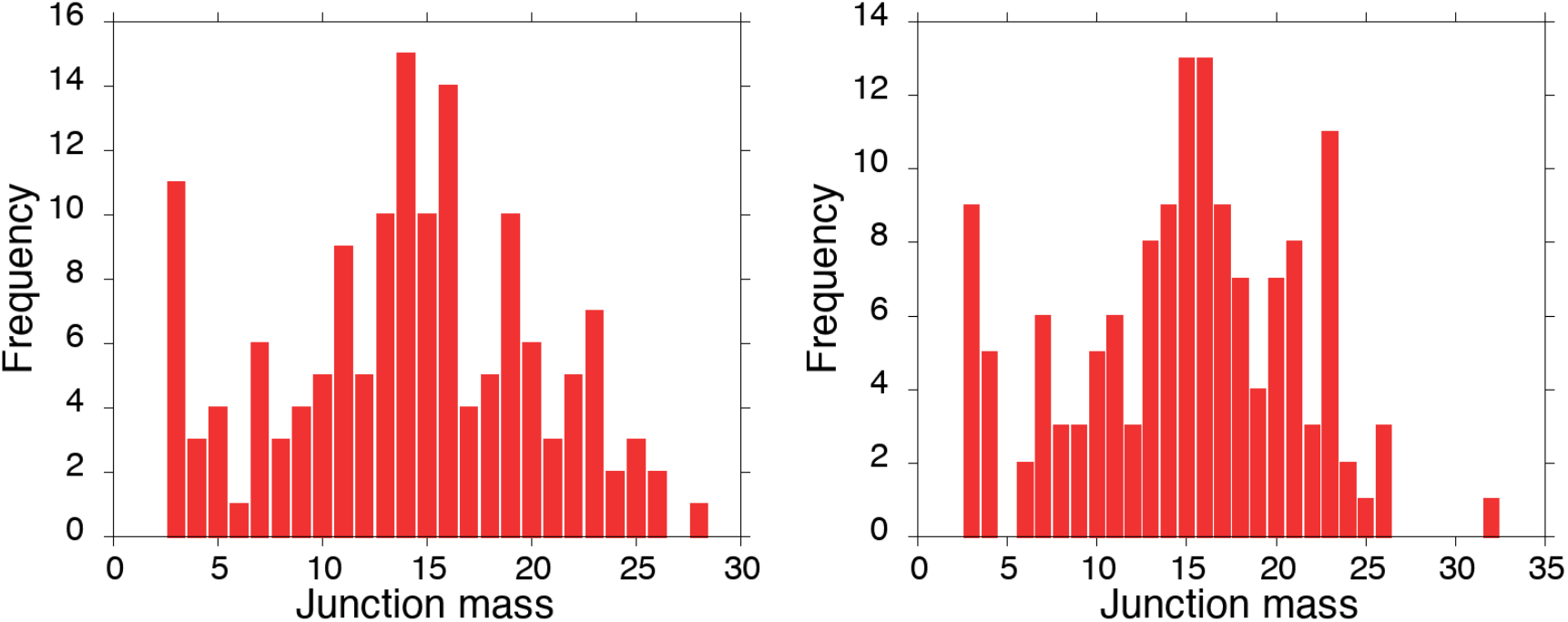
A) Histograms of the relative frequency of the number of polymers meeting at a junction (the junction *mass*) for an equilibrated network of 1215 B_16_ polymers with high affinity (ε = 0.8) (left) and the same network 500,000 time-steps later (right). The mass distribution is broad and approximately normal with a mean value around 15 polymers/junction. The enhanced peak for junctions with 3-4 polymers is likely due to junctions on the surface of the network being largely surrounded by solvent compared to those in the network interior. The variation in the histograms shows that polymers redistribute among the junctions over time reflecting the network’s fluid state (see also Figure S5).

By contrast, Figure 8B shows that the junction mass distribution for the same B_16_ polymers with low-affinity end-caps is exponential and rarely has more than 5 polymers/junction except at very high concentrations. The weak affinity precludes many polymers binding at the network junctions, but their mean mass is again invariant over time as seen from similarity of the histograms at different times.

**Fig. 8.**
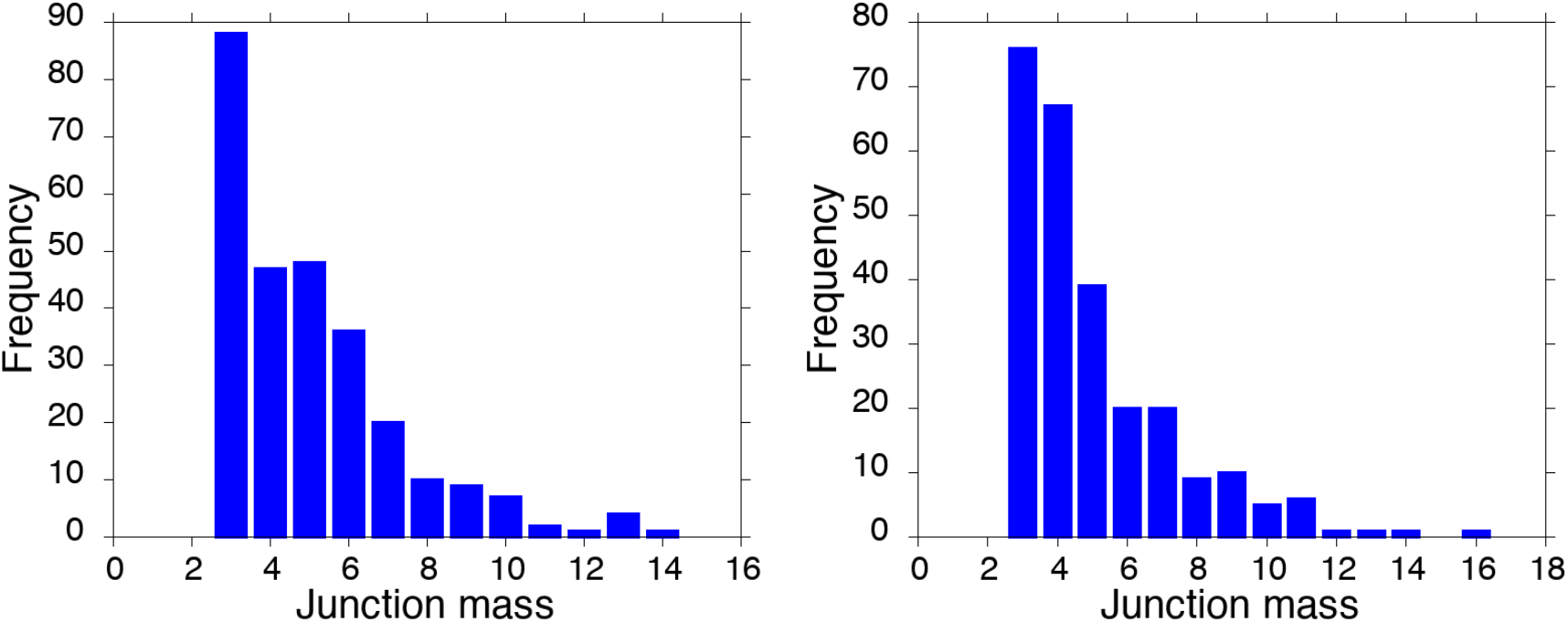
B) Histograms of the relative frequency of the junction mass for an equilibrated network of 1215 B_16_ polymers with low affinity (ε = 0.68) (left) and the same network 500,000 time-steps later (right). In contrast to the high-affinity case, the distribution is exponential with few junctions having more than 10 polymers. The variability in the histograms reflects the fluid state of the network.

## IV. CONCLUSIONS

In this work, we have elucidated the phase behavior and internal structure of a model biomolecular condensate and their dependence on the constituent molecules’ length and binding affinity using coarse-grained DPD simulations. Our results show that liquid-liquid phase separation of model IDPs has a richer behavior than has been reported in the literature, and does not require hydrophobic regions in the molecules.

The IDPs are represented as semi-flexible, hydrophilic polymers with sticky endcaps in a good solvent. In certain regimes of backbone length and end-cap affinity, the polymers phase separate at very low concentrations into a dense fluid droplet within a dilute phase. Although the condensed phase has a much greater polymer concentration than the surrounding dilute phase, it is still very porous, and the polymers retain large conformational fluctuations as seen experimentally.^13^ This phase separation is distinct from that reported in the simulations of Statt *et al*.^39^ as it occurs in the absence of hydrophobic regions in the polymers. The phase separation disappears below a critical, polymer length-dependent, value of the affinity, whereupon the polymers continuously merge into box-spanning networks on increasing concentration in agreement with Monte Carlo simulations in the literature.^38^

Biomolecular condensates are believed to be organizing centers for cellular biochemistry.^2,3^ Their porosity influences the diffusion of, and therefore the reactions between, proteins within them. Because it is hard for cells to control the size of phase separated droplets, particularly close to a critical point, it is important for their regulatory role that their internal structure be invariant to changes in their size. We find that the separation of connected junctions scales with the polymer backbone length as a self-avoiding random walk for both condensed droplets and box-spanning networks. This result is independent of the polymer concentration (and droplet morphology) over a wide range. It is also independent of the end-cap affinity (providing it is sufficiently large that the condensed phase exists) apart from a small affinity-dependent prefactor. This scaling behavior is distinct from that of hydrogels, for which the cross-link separation *decreases* with increasing concentration due to compression of the *permanently* cross-linked polymers.^55^ Adding polymers to the system causes the condensed phase to grow while the junction separation remains unchanged. On reaching the simulation box size, the junction separation remains constant and additional polymers add to the already-connected junctions. The robust structure we observe in the condensed phase provides a well-defined, stable, fluid environment within which biochemical reactions may be regulated. IDPs possess multiple, reversible interaction domains of various modalities,^16,18^ and our results support the idea that biomolecular condensates may regulate the interactions of client proteins by adjusting their internal porosity by modulating the separation between active binding sites on the scaffold or client proteins.

Our results also support the hypothesis that pathological fibrils form within a biomolecular condensate when the conformational dynamics of the constituent IDP’s are reduced.^56^ LLPS of the α-synuclein, an IDP associated with Parkinson’s disease, has been found to precede its aggregation into fibrils *in vitro* and in cells under oxidative stress.^57^ In a healthy state, IDPs *reversibly* bind and retain strong local disorder,^20^ as seen in our simulations. Post-translational modification of IDPs such as tau,^58^ or interactions with chaperone proteins,^59^ modify their conformational ensemble.^60^ If additional binding sites are thereby brought within range, the IDPs may progressively bind more tightly^9,61^ and transform into rigid fibrils.^56^ Experiments are now testing whether perturbing the conformational ensemble of a disease-prone IDP by adding a weakly-interacting ligand has therapeutic potential.^62^ Further interventions being investigated include adding a disulphide bond to α-synuclein,^63^ or genetically modifying cells to produce IDPs with alternative PTM sites.^64^ Opto-genetic experiments^65^ that add an inert linker to an IDP may also modify their phase behaviour. Although our model is a greatly-simplified representation of IDPs, it demonstrates the minimal molecular features that are sufficient to observe liquid-liquid phase separation of soluble, telechelic polymers. The model’s phase diagram may have additional regions yet to be explored. Greater fidelity with biological IDPs could be achieved by placing multiple binding sites on the polymers, and they could have a location-dependent backbone stiffness. Coarse-grained simulations can stimulate progress in this field by mapping out the consequences of experimental modifications of IDPs on their phase diagram and the material properties of biomolecular condensates.

## SUPPLEMENTARY MATERIAL

Technical details of the analysis can be found in the associated **supplementary material**. Snapshots and movies were produced using the open-source VMD software from the University of Illinois Urbana Champaign (http://www.ks.uiuc.edu/Research/vmd/).^66^ The analysis was performed using custom python code written by the authors, and included algorithms from the open source library scikit-learn (https://scikit-learn.org/stable/index.html) as described in the supplementary material.

The Supplementary Material includes the following movies:

Movie 1 Initial stage of the aggregation of 634 polymers B_16_ with low affinity ε = 0.68.
Movie 2 Initial stage of the aggregation of 634 polymers B_16_ with high affinity ε = 0.8.
Movie 3 Initial stage of the aggregation of 634 polymers B_16_ with very high affinity ε = 0.96.
Movie 4 Initial stage of the aggregation of 1251 polymers B_8_ with high affinity ε = 0.8.
Movie 5 Initial stage of the aggregation of 1180 polymers B_24_ with high affinity ε = 0.8.

## DATA AVAILABILITY STATEMENT

The executable DPD simulation code and the data that support the findings of this study are available from the corresponding author on reasonable request.

## ACKNOWLEDGEMENTS

The authors express their gratitude to W. Pezeshkian for many interesting discussions and for the script to export simulation snapshot files in VMD format. This study was supported by funding to the Blue Brain Project, a research centre of the École polytechnique fédérale de Lausanne (EPFL), from the Swiss government’s ETH Board of the Swiss Federal Institutes of Technology. The authors gratefully acknowledge computer time provided by the Blue Brain Project and Swiss National Supercomputing Centre and the Abacus 2.0 Super-Computing cluster at the University of Southern Denmark.

## AUTHOR CONTRIBUTIONS

JCS and JHI conceived the study; JCS performed the simulations; MB and EC wrote the analysis code. MB, EC, and JCS performed the data analysis, and all authors discussed the results; JCS and JHI wrote the manuscript and all authors commented on and revised the manuscript.

## Notes

### Competing Interest Statement

The authors have declared no competing interest.

## REFERENCES

1 S. Boeynaems, S. Alberti, N. L. Fawzi, T. Mittag, M. Polymenidou, F. Rousseau, J. Schymkowitz, J. Shorter, B. Wolozin, L. Van Den Bosch, P. Tompa, and M. Fuxreiter, “Protein Phase Separation: A New Phase in Cell Biology,” Trends Cell Biology 28, 420–435 (2018).

2 S. F. Banani, H. O. Lee, A. A. Hyman, and M. K. Rosen, “Biomolecular Condensates: Organizers of Cellular Biochemistry,” Nature Rev. Mol. Cell Biol. 18, 285–298 (2017).

3 A. S. Holehouse and R. V. Pappu, “Functional Implications of Intracellular Phase Transitions,” Biochemistry 57, 2415–2423 (2018).

4 A. Patel, H. O. Lee, L. Jawerth, S. Maharana, M. Jahnel, M. Y. Hein, S. Stoynov, J. Mahamid, S. Saha, T. M. Franzmann, A. Pozniakovski, I. Poser, N. Maghelli, L. A. Royer, M. Weigert, E. W. Myers, S. Grill, D. Drechsel, A. A. Hyman, and S. Alberti, “A Liquid-to-Solid Phase Transition of the ALS Protein FUS Accelerated By Disease Mutation,” Cell 162, 1066–1077 (2015).

5 P. A. Chong and J. D. Forman-Kay, “Liquid-Liquid Phase Separation in Cellular Signalling Systems,” Curr. Op. Struct. Biol. 41, 180–186 (2016);

5a P. Li, S. Banjade, H-C. Cheng, S. Kim, B. Chen, L. Guo, M. Llaguno, J. V. Hollingsworth, D. S. King, S. F. Banani, P. S. Russo, Q-X. Jiang, B. T. Nixon, and M. K. Rosen, “Phase Transitions in the Assembly of Multivalent Signalling Proteins,” Nature 483, 336–341 (2012);

5b X. Su, J. A. Ditlev, E. Hui, W. Xing, S. Banjade, J. Okrut, D. S. King, J. Taunton, M. K. Rosen, and R. D. Vale, “Phase Separation of Signaling Molecules Promotes T Cell Receptor Signal Transduction,” Science 352, 595–599 (2016).

6 M. Zeng, Y. Shang, Y. Araki, T. Guo, R. L. Huganir, and M. Zhang, “Phase Transition in Postsynaptic Densities Underlies Formation of Synaptic Complexes and Synaptic Plasticity,” Cell 166, 1163–1175 (2016).

7 D. Milovanovic, Y. Wu, X. Bian, and P. De Camilli, “A Liquid Phase of Synapsin and Lipid Vesicles,” Science 361, 604–607 (2018).

8 Y. Shin, Y-C. Chang, D. S. W. Lee, J. Berry, D. W. Sanders, P. Ronceray, N. S. Wingreen, M. Haataja, and C. P. Brangwynne, “Liquid Nuclear Condensates Mechanically Sense and Restructure the Genome,” Cell 175, 1481–1491 (2018).

9 S. Alberti and A. A. Hyman, “Are Aberrant Phase Transitions a Driver of Cellular Aging?,” Bioessays 38, 959–968 (2016).

10 A. Aguzzi and M. Altmeyer, “Phase Separation: Linking Cellular Compartmentalization to Disease,” Trends Cell Biology 26, 547–558 (2016).

11 S. Alberti, A. Gladfelter, and T. Mittag, “Considerations and Challenges in Studying Liquid-Liquid Phase Separation and Biomolecular Condensates,” Cell 176, 419–434 (2019).

12 D. Bracha, M. T. Walls, and C. P. Brangwynne, “Probing and engineering liquid-phase organelles,” Nat Biotechnol 37 (12), 1435–1445 (2019).

13 A. C. Murthy, G. L. Dignon, Y. Kan, G. H. Zerze, S. H. Parekh, J. Mittal, and N. L. Fawzi, “Molecular Interactions Underlying Liquid-Liquid Phase Separation of the FUS Low-Complexity Domain,” Nature Struct. Mol. Biol. 26, 637–648 (2019).

14 D. S. W. Protter, B. S. Rao, B. V. Treeck, Y. Lin, L. Mizoue, M. K. Rosen, and R. Parker, “Intrinsically Disordered Regions Can Contribute Promiscuous Interactions to RNP Granule Assembly,” Cell Reports 22, 1401–1412 (2018).

15 C. J. Brown, A. K. Johnson, A. K. Dunker, and G. W. Daughdrill, “Evolution and Disorder,” Current Opinion in Structural Biology 21, 441–446 (2011);

18. C. J. Oldfield and A. K. Dunker, “Intrinsically Disordered Proteins and Intrinsically Disordered Protein Regions,” Annu. Rev. Biochem. 83, 553–584 (2014).

16 C. P. Brangwynne, P. Tompa, and R. V. Pappu, “Polymer Physics of Intracellular Phase Transitions,” Nature Physics 11, 899–904 (2015).

17 A. S. Holehouse and R. V. Pappu, “Collapse Transitions of Proteins and the Interplay Among Backbone, Sidechain, and Solvent Interactions,” Annu. Rev. Biophys. 47, 19–39 (2018).

18 J. Wang, J-M. Choi, A. S. Holehouse, H. O. Lee, X. Zhang, M. Jahnel, S. Maharana, R. Lemaitre, A. Pozniakovsky, D. Drechsel, I. Poser, R. V. Pappu, S. Alberti, and A. A. Hyman, “A Molecular Grammar Governing the Driving Forces for Phase Separation of Prion-Like RNA Binding Proteins,” Cell 174, 688–699 (2018).

19 D. M. Mitrea, B. Chandra, M. C. Ferrolino, E. B. Gibbs, M. Tolbert, M. R. White, and R. W. Kriwacki, “Methods for Physical Characterization of Phase-Separated Bodies and Membraneless Organelles,” J. Mol. Biol. 430, 4773–4805 (2018).

20 K. A. Burke, A. M. Janke, C. L. Rhine, and N. L. Fawzi, “Residue-by-Residue View of In Vitro FUS Granules that Bind the C-Terminal Domain of RNA Polymerase II,” Molecular Cell 60, 231–241 (2015).

21 M-T. Wei, S. Elbaum-Garfinkle, A. S. Holehouse, C. C-H. Chen, M. Feric, C. B. Arnold, R. D. Priestley, R. V. Pappu, and C. P. Brangwynne, “Phase Behaviour of Disordered Proteins Underlying Low Density and High Permeability of Liquid Organelles,” Nature Chemistry 9, 1118–1125 (2017).

22 T. M. Franzmann, M. Jahnel, A. Pozniakovsky, J. Mahamid, A. S. Holehouse, E. Nüske, D. Richter, W. Baumeister, S. W. Grill, R. V. Pappu, A. A. Hyman, and S. Alberti, “Phase Separation of a Yeast Prion Protein Promotes Cellular Fitness,” Science 359, eaao5654-5651–eaao5654-5658 (2018).

23 Y. Wang, A. Lomakin, S. Kanai, R. Alex, and G. B. Benedek, “Liquid-Liquid Phase Separation in Oligomeric Peptide Solutions,” Langmuir 33, 7715–7721 (2017).

24 S. F. Banani, A. M. Rice, W. B. Peeples, Y. Lin, S. Jain, R. Parker, and M. K. Rosen, “Compositional Control of Phase-Separated Cellular Bodies,” Cell 166, 651–663 (2016).

25 M. Rubinstein and R. H. Colby, Polymer Physics. (Oxford University Press, New York, 2003).

26 K. M. Ruff, R. V. Pappu, and A. S. Holehouse, “Conformational Preferences and Phase Behaviour of Intrinsically Disordered Low Complexity Sequences: Insights from Multiscale Simulations,” Curr. Op. Struct. Biol. 56, 1–10 (2019);

26a G. L. Dignon, W. Zheng, and J. Mittal, “Simulation methods for liquid-liquid phase separation of disorded proteins,” Curr. Op. Chem. Eng. 23, 92–98 (2019).

27 S. Rauscher, V. Gapsys, M. J. Gajda, M. Zweckstetter, B. L. de Groot, and H. Grubmüller, “Structural Ensembles of Intrinsically Disordered Proteins Depend Strongly on Force Field: A Comparison to Experiment,” J. Chem. Theory and Comp. 11, 5513–5524 (2015).

28 H. Kang, B. Luan, and R. Zhou, “Glassy Dynamics in Mutant Huntingtin Proteins,” J. Chem. Phys. 149, 072333–072341 (2018).

29 A. Balupuri, K-E. Choi, and N. S. Kang, “Computational Insights into the Role of α-Strand/Sheet in Aggregation of α-Synuclein,” Scientific Reports 9, 59-51–59-13 (2019).

30 A. J. Lyons, N. S. Gandhi, and R. L. Mancera, “Molecular Dynamics Simulation of the Phosphorylation-Induced Conformational Changes of a Tau Peptide Fragment,” Proteins 82, 1907–1923 (2014).

31 B. Barz, Q. Liao, and B. Strodel, “Pathways of Amyloid-β Aggregation Depend on Oligomer Shape,” JACS 140, 319–327 (2017).

32 G. L. Dignon, W. Zheng, Y. C. Kim, R. B. Best, and J. Mittal, “Sequence Determinants of Protein Phase Behavior from a Coarse-Grained Model,” PLoS Computational Biology 14, e1005941 (2018).

33 T. S. Harmon, A. S. Holehouse, and R. V. Pappu, “Differential Solvation of Intrinsically Disordered Linkers Drives the Formation of Spatially Organised Droplets in Ternary Systems of Linear Multivalent Proteins,” New. J. Physics 20, 045002–045016 (2018).

34 A. Chatteraj, M. Youngstrom, and L. M. Loew, “The Interplay of Structural and Cellular Biophysics Controls Clustering of Multivalent Molecules,” Biophys. J. 116, 560–572 (2019).

35 N. Arai, “Structural Analysis of Telechelic Polymer Solution Using Dissipative Particle Dynamics Simulations,” Mol. Sim. 41, 996–1001 (2015).

36 C. Xiao and D. M. Heyes, “Brownian Dynamics Simulations of Attractive Polymers in Solution,” J. Chem. Phys. 117, 2377–2388 (2002).

37 C. Manassero, G. Raos, and G. Allegra, “Structure of Model Telechelic Polymer Melts by Computer Simulation,” J. Macromol. Sci. Part B: Physics 44, 855–871 (2005).

38 T. S. Harmon, A. S. Holehouse, M. K. Rosen, and R. V. Pappu, “Intrinsically disordered linkers determine the interplay between phase separation and gelation in multivalent proteins,” eLife 6, e30294 (2017).

39 A. Statt, H. Casademunt, C. P. Brangwynne, and A. Z. Panagiotopoulos, “Model for disordered proteins with strongly sequence-dependent liquid phase behaviour,” J. Chem. Phys. 152, 075101 (2020).

40 P. J. Hoogerbrugge and J. M. V. A. Koelman, “Simulating Microscopic Hydrodynamic Phenomena with Dissipative Particle Dynamics,” Europhys. Lett. 19, 155–160 (1992);

40a P. Espagnol and P. B. Warren, “Statistical Mechanics of Dissipative Particle Dynamics,” Europhysics Letters 30, 191–196 (1995).

41 R. D. Groot and P. B. Warren, “Dissipative Particle Dynamics: Bridging the Gap Between Atomistic and Mesoscopic Simulations,” J. Chem. Phys. 107, 4423–4435 (1997).

42 A. A. Hyman and C. P. Brangwynne, “Beyond Stereospecificity: Liquids and Mesoscale Organization of Cytoplasm,” Dev. Cell 21, 14–16 (2011).

43 J-G. Gai, H-L. Li, C. Schrauwen, and G-H. Hu, “Dissipative Particle Dynamics Study on the Phase Morphologies of the Ultrahigh Molecular Weight Polyethylene/Polypropylene/Poly(ethylene glycol) Blends,” Polymer 50, 336–346 (2009);

43a H. Droghetti, I. Pagonabarraga, P. Carbone, P. Asinari, and D. Marchisio, “Dissipative Particle Dynamics Simulations of Tri-Block CoPolymer and Water: Phase Diagram Validation and Microstructure Identification,” J. Chem. Phys. 149, 184903–184913 (2018).

44 A. A. Gavrilov, Y. V. Kudryavtsev, and P. G. Khalatur, A. V. Chertovich, “Microphase Separation in Regular and Random Copolymer Melts by DPD Simulations,” Chem. Phys. Lett. 503, 277–282 (2011).

45 J. C. Shillcock and R. Lipowsky, “Equilibrium Structure and Lateral Stress Distribution of Amphiphilic Bilayers from Dissipative Particle Dynamics Simulations,” J. Chem. Phys. 117, 5048–5061 (2002).

46 L. Gao, J. Shillcock, and R. Lipowsky, “Improved dissipative particle dynamics simulations of lipid bilayers,” J Chem Phys 126 (1), 015101 (2007).

47 M. Laradji and P. B. Sunil Kumar, “Dynamics of Domain Growth in Self-Assembled Fluid Vesicles,” Phys. Rev. Lett. 93, 198105–198108 (2004).

48 J. C. Shillcock and R. Lipowsky, “Tension-induced fusion of bilayer membranes and vesicles,” Nat Mater 4 (3), 225–228 (2005);

48a A. Grafmuller, J. Shillcock, and R. Lipowsky, “The fusion of membranes and vesicles: pathway and energy barriers from dissipative particle dynamics,” Biophys J 96 (7), 2658–2675 (2009).

49 J. C. Shillcock, “Spontaneous Vesicle Self-Assembly: A Mesoscopic View of Membrane Dynamics,” Langmuir 28, 541–547 (2012).

50 K. A. Smith, D. Jasnow, and A. C. Balazs, “Designing Synthetic Vesicles that Engulf Nanoscopic Particles,” J. Chem. Phys. 127, 084703–084713 (2007);

50a W. Pezeshkian, H. Gao, S. Arumugam, U. Becken, P. Bassereau, J. C. Florent, J. H. Ipsen, L. Johannes, and J. C. Shillcock, “Mechanism of Shiga Toxin Clustering on Membranes,” ACS Nano 11 (1), 314–324 (2017).

51 M. Venturoli, M. M. Sperotto, M. Kranenburg, and B. Smit, “Mesoscopic Models of Biological Membranes,” Physics Reports 437, 1–54 (2006).

52 P. Espagnol and P. B. Warren, “Perspective: Dissipative Particle Dynamics,” J. Chem. Phys. 146, 150901–150917 (2017).

53 N. O. Taylor, M-T. Wei, H. A. Stone, and C. P. Brangwynne, “Quantifying Dynamics in Phase-Separated Condensates Using Fluorescence Recovery After Photobleaching,” Biophys. J. 117, 1285–1300 (2019).

54 A. N. Semenov, I. A. Nyrkova, and M. E. Cates, “Phase Equilibria in Solutions of Associating Telechelic Polymers: Rings vs Reversible Network,” Macromolecules 28, 7879–7885 (1995).

55 P. J. Flory, Principles of Polymer Chemistry. (Cornell University Press, Ithaca, 1953).

56 B. Tüu-Szabó, G. Hoffka, N. Duro, and M. Fuxreiter, “Altered dynamics may drift pathological fibrillization in membraneless organelles,” BBA - Proteins and Proteomics 1867, 988–998 (2019).

57 S. Ray, N. Singh, S. Pandey, R. Kumar, L. Gadhe, D. Datta, K. Patel, J. Mahato, Navalkar A, R. Panigrahi, D. Chatterjee, S. Maiti, S. Bhatia, S. Mehra, A. Singh, J. Gerez, A. Chowdhury, A. Kumar, R. Padinhateeri, R. Riek, G. Krishnamoorthy, and S. K. Maji, “Liquid-liquid phase separation and liquid-to-solid transition mediate alpha-synuclein amyloid fibril containing hydrogel formation,” bioRxiv preprint, 1–40, http://dx.doi.org/10.1101/619858 (2019).

58 S. Zhu, A. Shala, A. Bezginov, A. Sljoka, G. Audette, and D. J. Wilson, “Hyperphosphorylation of Intrinsically Disordered Tau Protein Induces an Amyloidogenic Shift in Its Conformational Ensemble,” PLoS One 10, e0120416 (2015).

59 Z. Wang and H. Zhang, “Phase Separation, Transition, and Autophagic Degradation of Proteins in Development and Pathogenesis,” Trends Cell Biology 29, 417–427 (2019).

60 E. E. Boczek and S. Alberti, “Phase changes in neurotransmission,” Science 361, 548–549 (2018).

61 T. R. Peskett, F. Rau, J. O’Driscoll, R. Patani, A. R. Lowe, and H. R. Saibil, “A Liquid To Solid Phase Transition Underlying Pathological Huntingtin Exon1 Aggregation,” Mol. Cell 70, 588–601 (2018).

62 B. K. Maity, V. Vishvakarma, D. Surendran, A. Rawat, A. Das, S. Pramanik, N. Arfin, and S. Maiti, “Spontaneous Fluctuations Can Guide Drug Design Strategies for Structurally Disordered Proteins,” Biochemistry 57, 4206–4213 (2018).

63 A. Carija, F. Pinheiro, J. Pujols, I. C. Brás, D. F. Lázaro, C. Santambrogio, R. Grandori, T. F. Outeira, S. Navarro, and S. Ventura, “Biasing the α-Synuclein Conformational Ensemble Towards Compact States Abolishes Aggregation and Neurotoxicity,” Redox Biology 22, 101135–101150 (2019).

64 Z. Monahan, V. H. Ryan, A. M. Janke, K. A. Burke, S. N. Rhoads, G. H. Zerze, R. O’Meally, G. L. Dignon, A. E. Conicella, W. Zheng, R. B. Best, R. N. Cole, J. Mittal, F. Shewmaker, and N. L. Fawzi, “Phosphorylation of the FUS Low-Complexity Domain Disrupts Phase Separation, Aggregation and Toxicity,” EMBO Journal 36, 2951–2967 (2017).

65 Y. Shin, J. Berry, N. Pannucci, M. P. Haataja, J. E. Toettcher, and C. P. Brangwynne, “Spatiotemporal Control of Intracellular Phase Transitions Using Light-Activated OptoDroplets,” Cell 168, 159–171 (2017).

66 W. Humphrey, A. Dalke, and K. Schulten, “VMD - Visual Molecular Dynamics,” Journal of Molecular Graphics 14, 33–38 (1996).

