## Supplementary Material for "Structure of biomolecular condensates from dissipative particle dynamics simulations"

#### 1) Relation between end-cap binding affinity and DPD conservative parameters

The conservative interaction parameter $a_{ij}$ between two different bead types i,j in DPD is related to their mutual solubility.[1] Because the simulated IDPs are hydrophilic polymers, they are soluble in the aqueous solvent. An attraction is created between the polymers' end-caps by setting $a_{EE} < a_{WW}$, which causes them to prefer associating with each other instead of being solvated. We define a dimensionless *affinity* in terms of the conservative interaction parameters between the end-cap beads and the solvent ($a_{WE}$) and the end-cap selfinteraction ($a_{EE}$) by:

$$\epsilon = (a_{WE} - a_{EE})/a_{WE} .$$

We have simulated the behavior of systems of polymers with a wide range of affinities. In the limit ε = 0, the IDPs reduce to non-associating, semi-flexible polymers. Polymers with affinities below ε = 0.68 were not observed to phase separate nor form a single, box-spanning network at high concentrations. Subsequent investigations were limited to affinities larger than this. For ease of discussion, we use the following *qualitative* labels to refer to affinities in distinct ranges: very strong affinity (ε ~ 0.96), strong (ε ~ 0.8), and weak (ε ~ 0.68).

**2) Algorithm for identifying the Largest Equilibrium Network**

At very low concentrations, the telechelic polymers are dispersed in the bulk solvent. As their concentration increases, they aggregate into transient clusters or a network. Observation of the simulations of polymers with low-affinity end-caps showed that a fraction of them remain dispersed in the bulk solvent even when a network has formed. In such cases, single polymers or small clusters merge and break up during the simulation. By contrast, only one or a few polymers with high affinity end-caps remain free in the bulk solvent. Because our goal is to relate the molecular properties of IDPs to the structure of biomolecular condensates, we want to study the largest available network as this is likely to be most similar to the micron-sized experimental systems. If several networks are present in the simulation, the question arises of which one, or set, of networks to use for calculating equilibrium properties. Small networks are unlikely to be structurally similar to experimental biomolecular condensates containing much larger numbers of proteins because of their large surface to volume ratio. In order to improve the accuracy of our results, we analyze the structural properties of only the largest network present at each sampling time during the simulation. We refer to this network as the *Largest Equilibrium Network* (LEN).

A clustering algorithm is used to identify all polymers that are connected by their end-caps into disconnected networks, and the largest such network is identified as the LEN. The LEN is *not* a static structure because polymers diffuse between junctions within the network and some detach and reattach during the simulation. But we expect that once the simulation has reached equilibrium, the largest network at each time best represents the experimental condensates that exchange components with the bulk phase. The LEN is recomputed for each sample taken from the simulations.

The clustering algorithm is DBSCAN from the open source scikit library (https://scikit-learn.org/stable/index.html). The algorithm sorts a set of spatially-distributed points into *clusters* based on their proximity and was chosen because it does not require the number of clusters to be specified *a priori*. and it allows noise, *i.e.*, single points or small clusters, to be excluded from analysis. Two parameters are needed to perform the clustering analysis: 1) the maximum separation $d_{max}$ between two points in space for them to be assigned to the same cluster; 2) the minimum number of points in a cluster. To reduce the number of points that have to be analyzed in constructing the LEN, each end-cap was represented by the central E bead directly connected to the polymer backbone. This reduces the computational load by a factor of 4. Two polymers are assigned to the same cluster (referred to as *junctions* hereafter and in the text) if their central end-cap beads are closer than a maximum separation $d_{max} = 1.5\ d_0$ in space. The value of $d_{max}$ was varied either side of this value but this resulted in no significant change to the LEN size. Once all the points have been classified, the LEN is identified as the largest connected set of junctions. Finally, junctions in the LEN whose size is less than $N_{min} = 3$ are discarded. This is to remove the distorting effect on the LEN properties of polymers with one dangling end, or junctions with only two polymers.

**3) Ring conformations of polymers with strong end-cap affinity**

Telechelic polymers with weak or no end-cap affinity behave as semi-flexible polymers in a good solvent. But polymers with stronger affinites are found to adopt ring conformations in which both their end-caps bind to each other. These conformations occur even at very low concentrations, and their proportion increases with increasing affinity. Figure S2 shows that polymers also spontaneously aggregate into small micelle-like structures. We do not analyze these small micelle-like structures further as we are interested in the equilibrium properties of the condensed network. But we take account of ring-like polymers when they occur within the LEN as described next.

A different definition of a ring conformation is used for polymers loose in the bulk solvent and those in the network. A polymer in the bulk solvent forms a ring if its end-caps are within a fixed distance 1.5 $d_0$ of each other. But a polymer in the network is in a ring conformation if both of its end-caps are attached to the same junction regardless of their spatial separation. A small fraction of the polymers in the networks persist in ring conformation at all concentrations studied, and are particularly common for high affinity polymers (Fig. S4). Polymers in ring-like conformations within the network can influence its structure. Rings have an-end-to-end length close to zero and are excluded from the calculation of the mean junction separation (Figs. 4 and 5). Polymers in the network that have one dangling end are also not counted in this calculation. However, polymers in ring conformations occupy space and may interact sterically with nearby polymers. Therefore, they are included in the calculation of the network size (Fig. 3) and the distribution of polymers among the junctions(Figs. 6 and 7).

**4) Statistical errors in structural properties of the network**

Statistical errors are estimated as follows. Simulations of each system with a fixed polymer architecture, affinity and concentration are carried out as a sequence of runs where each run is restarted from the final configuration of the previous run. Typically, 9 runs are performed of 600,000 steps each and the first

5 discarded so that the systems evolve for at least $3.10^6$ time-steps after their random initial state to allow them to reach equilibrium. Time-averaged observables, e.g., the network's junction separation and mean junction mass, are then sampled at 50,000 time-step intervals and averaged over several 600,000 time-step runs. Four such averages over successive 600,000 step intervals for the full range of polymer densities for high and low affinity are shown in Figures S7 and S8. The agreement between the curves shows that the networks are in equilibrium for both low and high affinity end-caps at all polymer concentrations studied except the lowest below 0.001. At these low densities, the (small) networks break up and reform continually leading to large fluctuations in its properties. Each run of 600,000 steps requires 5 cpu-days on a single Intel E5-2680v3 core, and therefore a complete sequence of 9 runs requires 45 cpu-days.

**5) System size dependence of network properties**

The networks we observe range from small spherical droplets to large structures that span the periodic boundaries of the simulation box. We have investigated whether the network's structural properties are affected by the box size by performing some simulations in a box with linear dimension $L = 64d_0$. Figure S9 shows that the mean junction separation is identical within statistical errors in the smaller and larger simulation boxes. The absence of any system size dependence further supports the conclusion that the self-assembled network is an equilibrium thermodynamic phase.

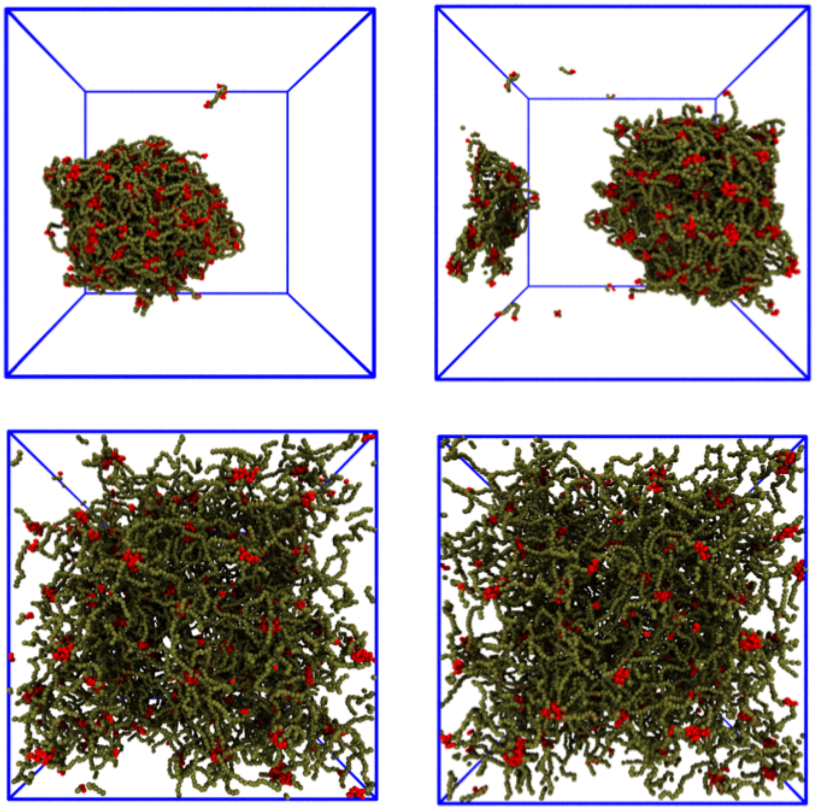

$B_8$ $B_{10}$

$B_{24}$ $B_{32}$

**Figure S1 Polymers of length $B_8$, $B_{10}$, $B_{24}$, $B_{32}$ aggregate into porous networks similar to those of $B_{16}$.** Polymers with very high affinity end-caps ($\varepsilon$ = 0.8) and backbone lengths of 8, 10, 24, and 32 beads show similar aggregation behavior to $B_{16}$ polymers (*cp.* bottom left snapshot in Fig. 2). Increasing the polymers' length at constant affinity changes the balance between their conformational entropy and binding affinity. This increases the spatial separation of the junctions and weakens their Note that there are 1251, 1242, 1180 and 891 polymers in the snapshots respectively. Disconnected pieces of polymer are connected via the periodic boundary conditions. Also note that solvent particles are always invisible in snapshots.

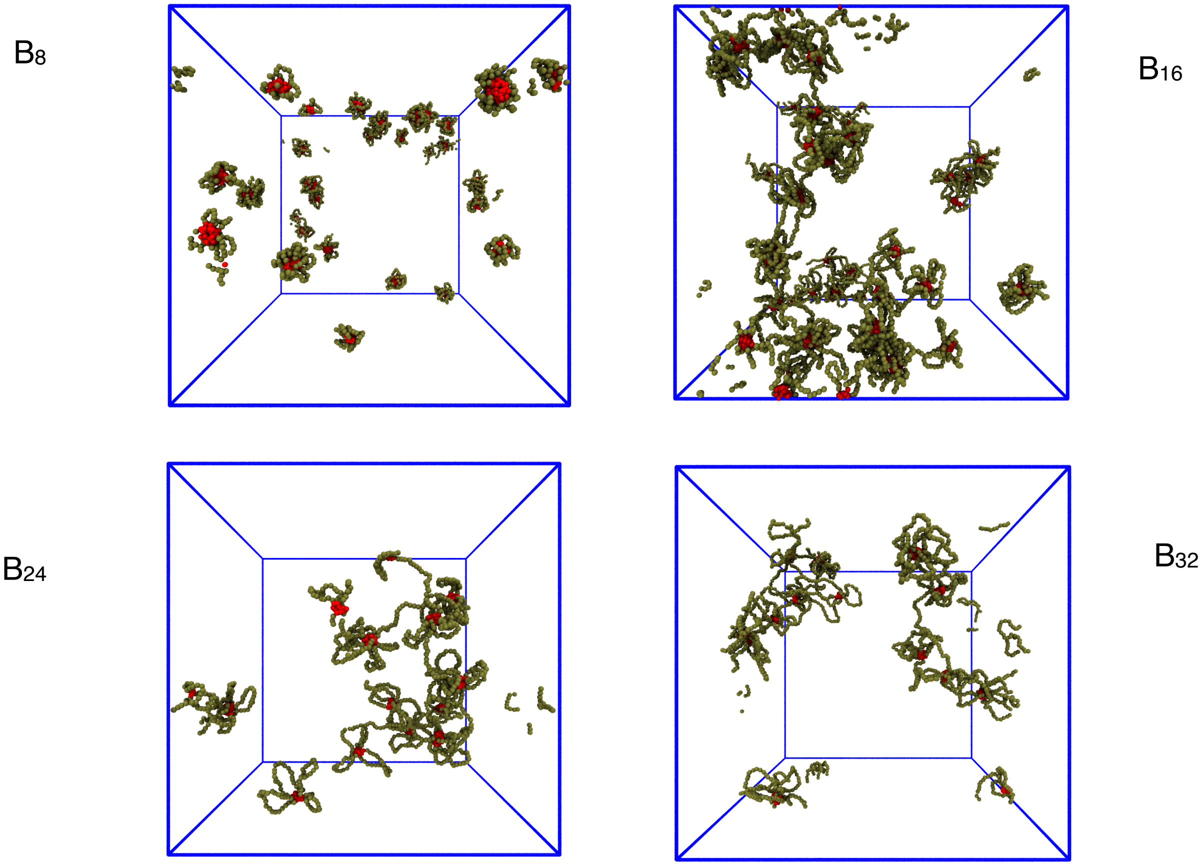


**Figure S2 Polymers with very high affinity (ε = 0.96) end-caps adopt ring-like structures in the dilute phase.** Short $B_8$ polymers (top left, 262 polymers) form small, micelle-like aggregates in the dilute phase driven by the strong affinity of the end-caps. A similar number of $B_{16}$ polymers (top right, 260 polymers) form a stringy network of connected clusters. Longer polymers $B_{24}$ (bottom left, 65 polymers) and $B_{32}$ (bottom right, 65 polymers) form stringy networks and "rosettes" at lower concentrations as the longer backbones allow the polymers to connect across larger distances. The high end-cap affinity stabilizes these structures and delays their aggregation into a large network.

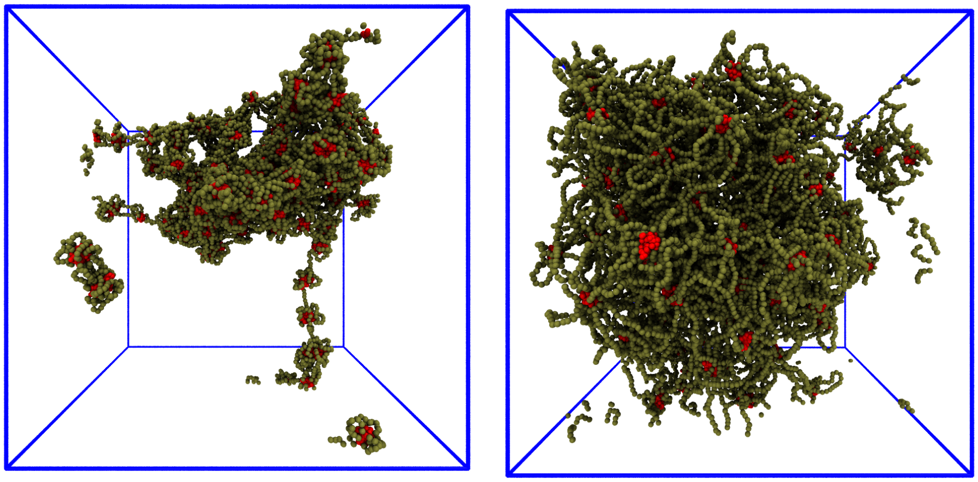

**Figure S3 Polymers with very high-affinity end-caps adopt ring-like conformations within the condensed network at high concentrations.** At high concentrations, polymers with high affinity (ε = 0.96) form a condensed network in which they adopt ring-like conformations with both end-caps meeting at the same junction. The proportion of rings is quantified in Fig. S4. For short polymers $B_8$ (left snapshot), the strong affinity transiently creates entropically unfavourable structures like the chain of rings in the lower half of the snapshot. Approximately one half of the $B_8$ polymers are in ring conformations at this concentration (0.004). Longer polymers, $B_{24}$ (right snapshot), at the same concentration form an extended porous network in which the fraction of rings is comparable to lower-affinity polymers. Fig. S4 shows that less than 20% of the $B_{24}$ polymers form rings in the network.

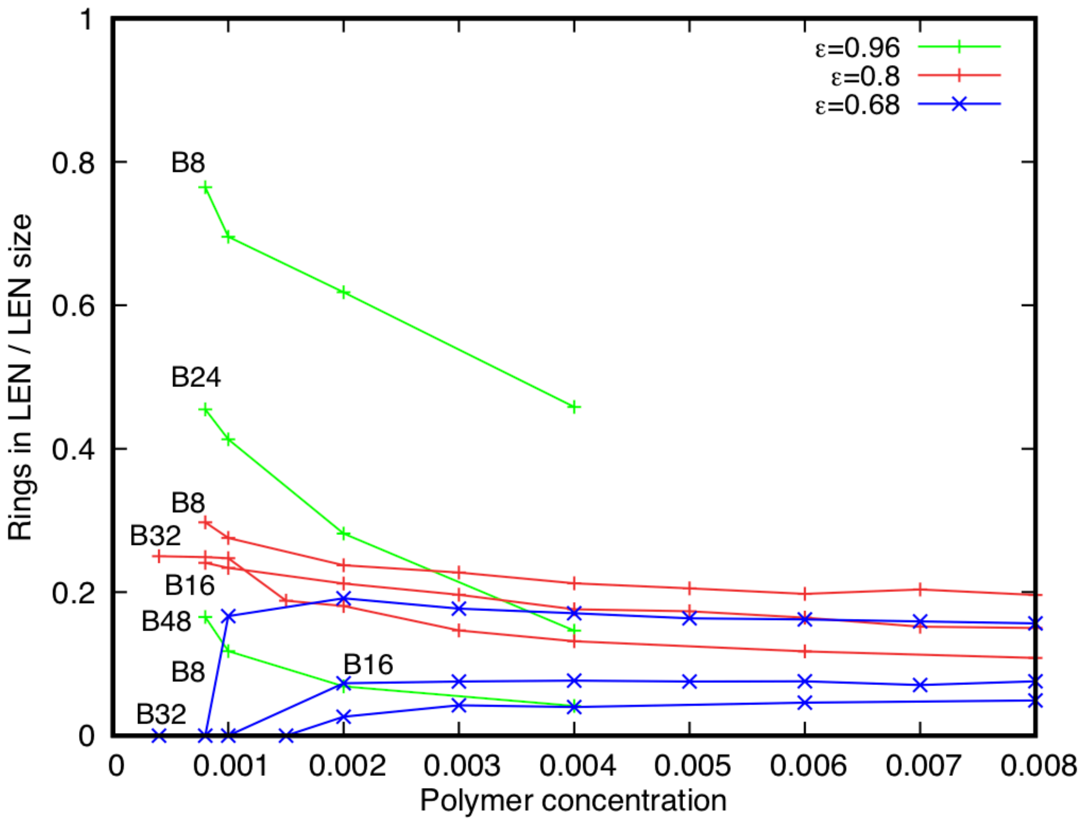


**Figure S4 Fraction of polymers in ring conformations within the LEN.** A polymer in a ring conformation in the LEN has both end-caps at the same junction. The number of polymers that form rings approaches a constant fraction (5 - 20%) of the LEN size at high concentrations, and this fraction has a weak dependence on the end-cap affinity and backbone length. Polymers with the highest affinity studied (ε = 0.96) form many ring conformations at low concentrations but the fraction falls to the range of 10-20% typical of polymers with weaker affinities. The curves for the highest affinity (ε = 0.96) only reach a concentration of 0.004, but the longer polymers ($B_{24}$, $B_{48}$) already show the same fraction of rings in the LEN as the lower-affinity polymers at this concentration.

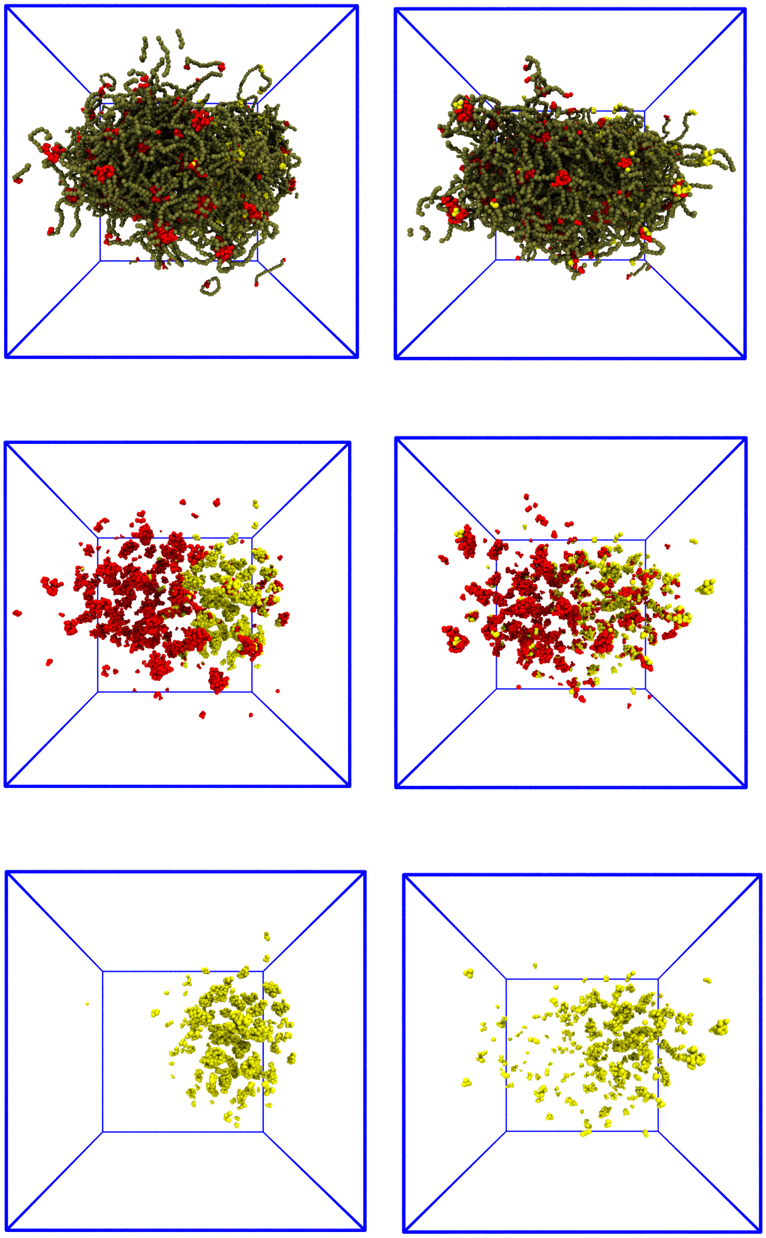

**Figure S5 Simulated Fluorescence Recovery after Photobleaching (FRAP) shows that the network phase is fluid.** The snapshots in the left-hand column show an equilibrated network of 1215 $B_{16}$ polymers with high affinity ($\varepsilon = 0.8$) while those in the right column show the same network 600,000 time-steps later. The top row shows all polymers; middle row - all polymer end-caps, bottom row -

*labelled* end-caps only. When the equilibrated simulation is restarted, the colour of the polymer end-caps located in the right-hand half of the network is changed from red to yellow to represent the bleaching effect in a FRAP experiment. The labelled polymers subsequently diffuse through the network as time passes indicating the fluid state of the network.

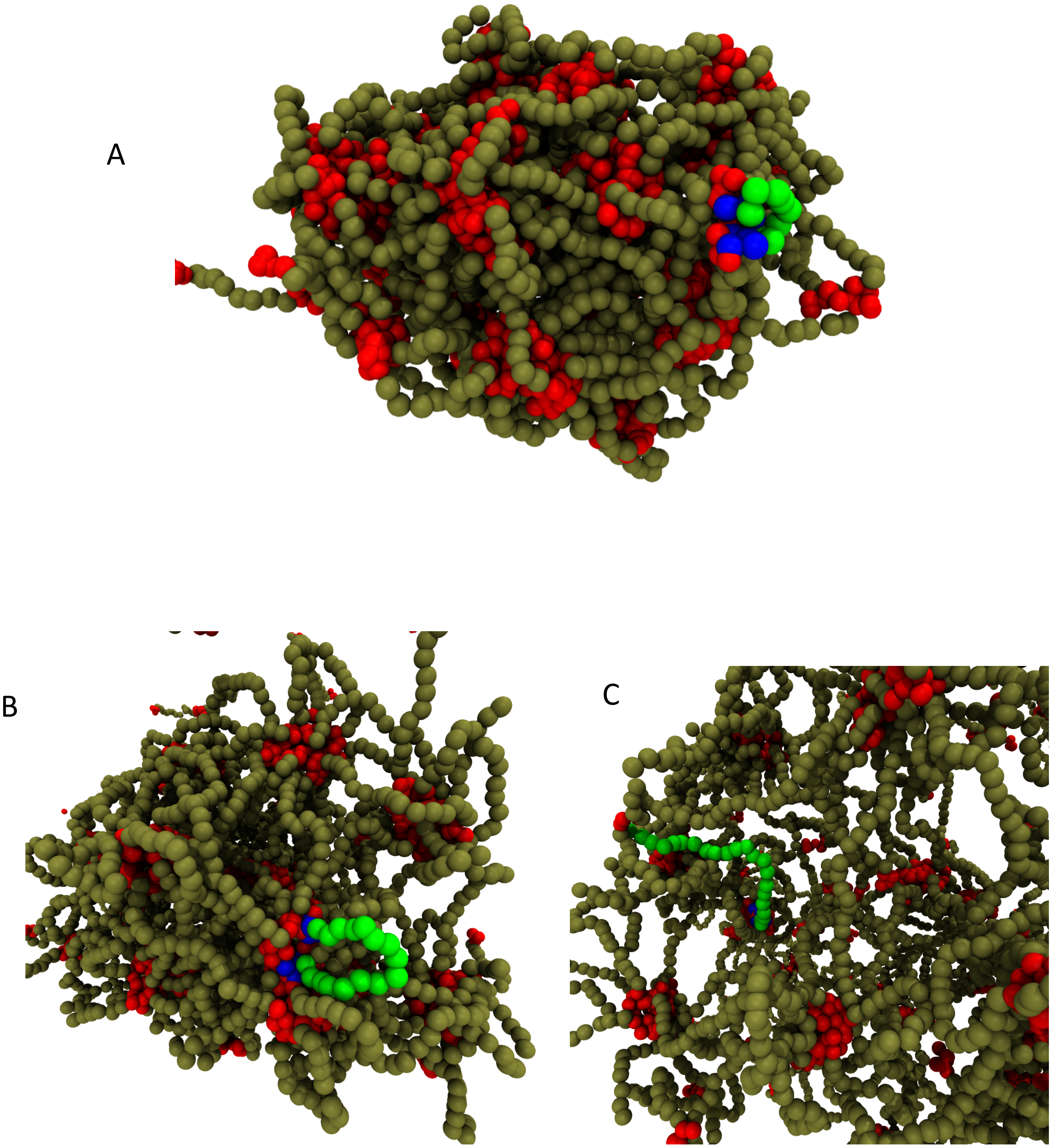


**Figure S6 Examples of single polymer conformations in the condensed phase for different backbone lengths.** A single polymer is selected randomly and colored with green backbone beads and blue end-caps. A) Short $B_8$

polymers with high affinity (ε = 0.8) often form tight rings at a junction; B) $B_{16}$ polymers with the same affinity form looser rings; C) $B_{24}$ polymers with the same affinity typically span spatially-separated junctions.  Note that the porosity of the networks increases with polymer backbone length. Fig. S4 shows that the proportion of polymers in ring conformations is below 20% for all polymer lengths at this end-cap affinity and concentrations.

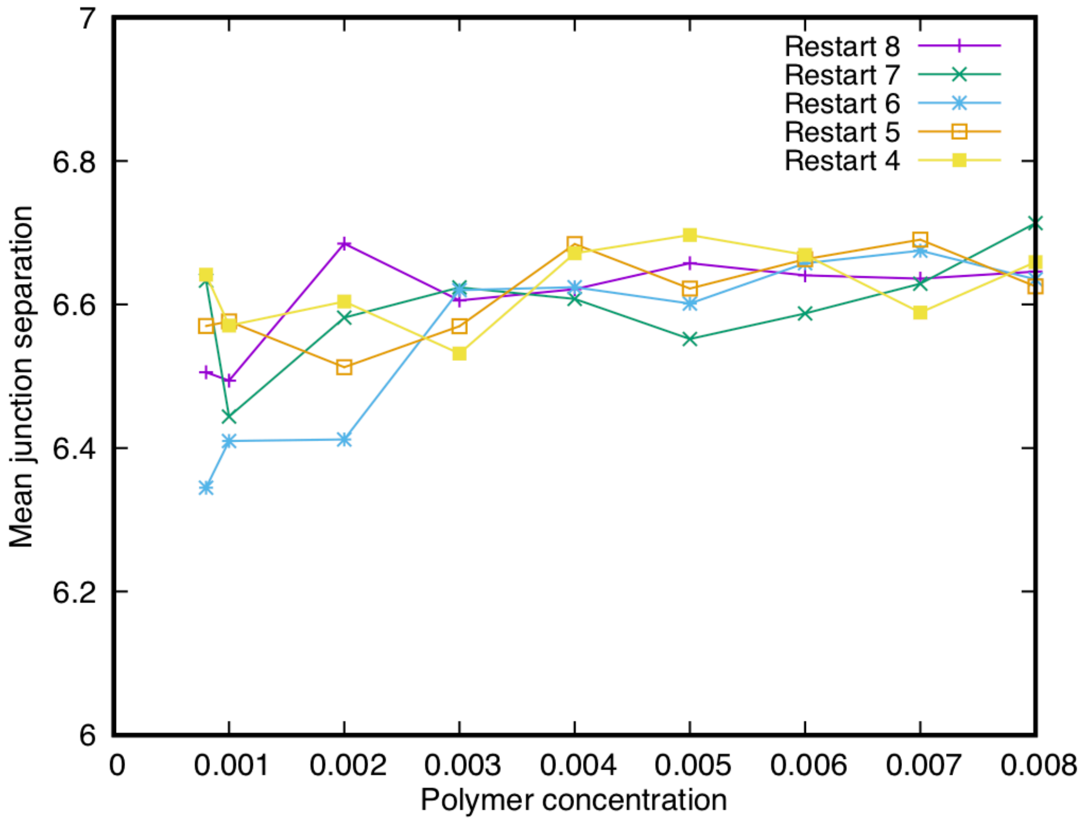


**Figure S7 The LEN of high affinity polymers is in equilibrium.** The curves show the mean junction separation in the LEN for $B_{16}$ polymers with *strong* affinity (ε = 0.8) as a function of the polymer concentration. Each curve is taken from a simulation restarted from the end of a previous one (Restart 4 - Restart 8). Each run was carried out for at least 300,000 time steps with the earliest curve (Restart 4) starting 1.2 million time steps after the simulation begins (hence, Restarts 1 - 3 were discarded). The mean junction separation shows small fluctuations about a stationary value at all concentrations showing that the LEN is in equilibrium.

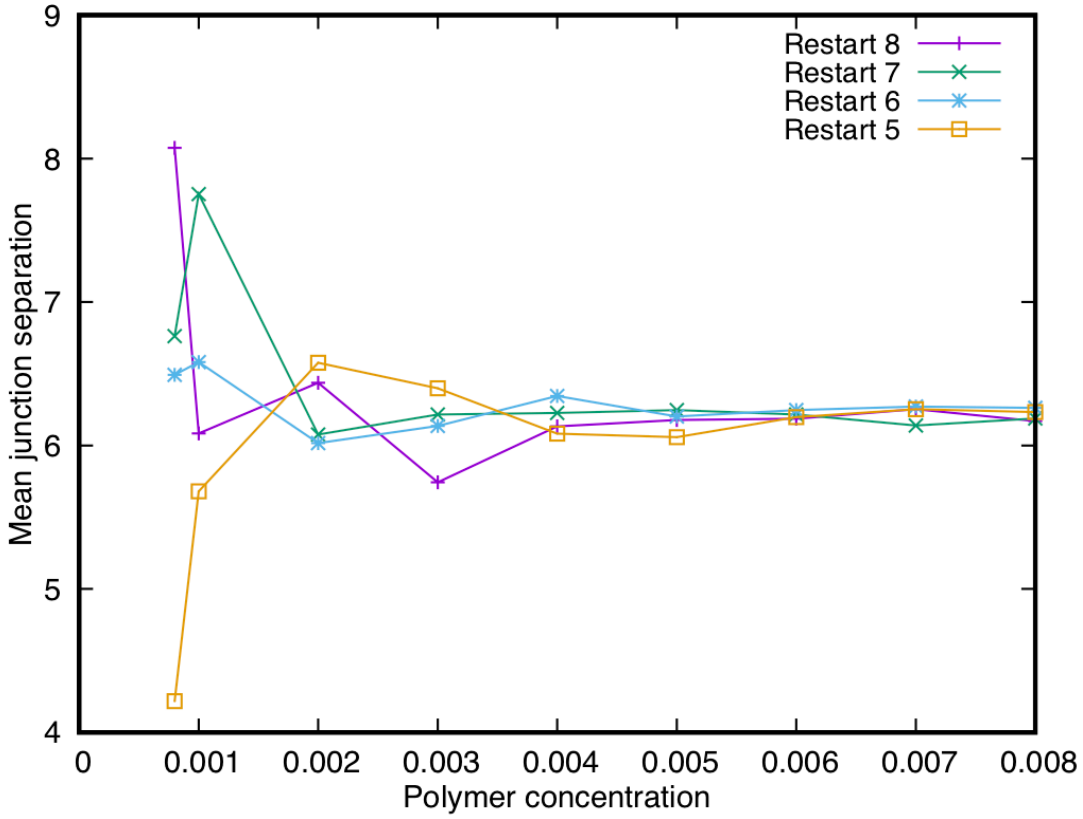


**Figure S8 The LEN of low-affinity polymers is in equilibrium.** Similar to Figure S7, the curves show the mean junction separation in the LEN for $B_{16}$ polymers with *weak* affinity (ε = 0.68) as a function of the polymer concentration. The curves are taken over four successive simulations (Restart 5 - Restart 8) containing at least 300,000 time steps each with the earliest curve (Restart 5) starting 1.5 million time steps after the simulation begins. Similar to Figure S7, the junction separation shows small fluctuations about a stationary mean value at all concentrations above 0.002 indicating that the LEN is in equilibrium. The large fluctuations at concentrations below 0.002 are due to the small size of the network and its instability for weak binding affinity.

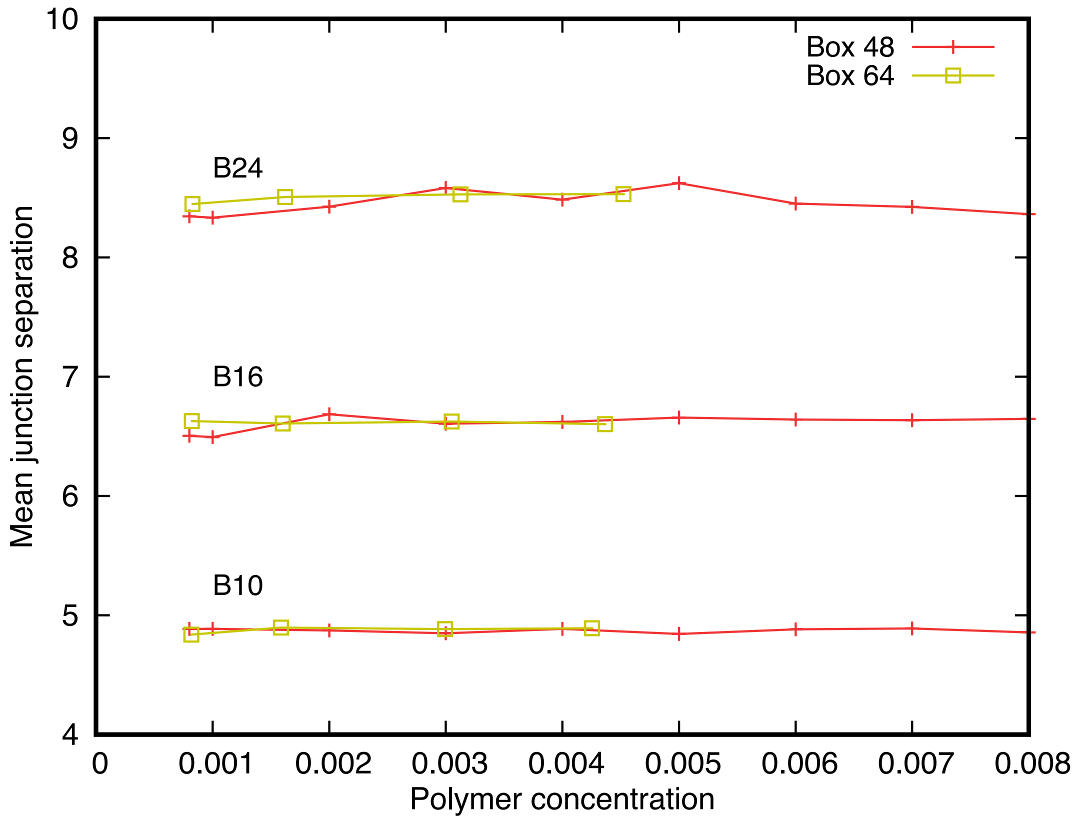


**Figure S9 System size dependence of the mean junction separation.** The mean junction separation in networks composed of polymers with a strong end-cap affinity (ε = 0.8) and backbone lengths of $B_{10}$, $B_{16}$, and $B_{24}$ is independent of the simulation box size. Snapshots of the networks in a simulation box $(48d_0)^3$ are shown in Figures 2 and S1, and snapshots of networks in the $(64d_0)^3$ box are shown in Figure S10.

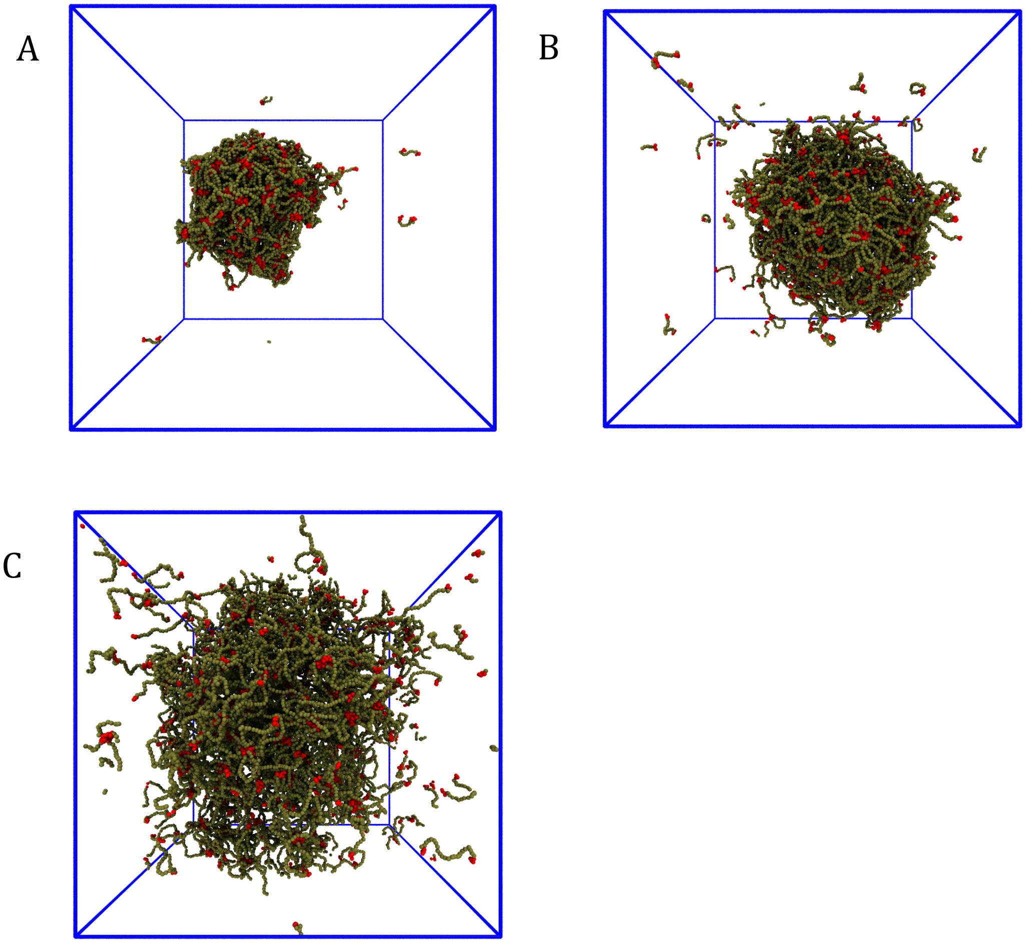


**Figure S10 Snapshots of networks in the larger simulation box $(64d_0)^3$.** The snapshots show networks of 1215 polymers with backbone lengths $B_{10}$, $B_{16}$, and $B_{24}$ and strong end-cap affinity ($\varepsilon$ = 0.8) in a simulation box $(64d_0)^3$. They should be compared with the corresponding snapshots in Figures 2 and S1 for a simulation box size $(48d_0)^3$. The network of $B_{10}$ polymers (A above and top right in Figure S1) forms a nearly-spherical droplet in both box sizes showing that the droplet is the equilibrium state. Networks of $B_{16}$ polymers show similar behavior (B above and bottom left in Figure 2), but Figure 2 has 634 $B_{16}$ polymers in

$(48d_0)^3$ while the above snapshot has 1215 $B_{16}$ polymers in $(64d_0)^3$. Also, the network in Figure 2 has connected to itself across the periodic boundaries of the simulation box which results in its elongated shape. The network of $B_{24}$ polymers (C above and bottom left in Figure S1) forms an extended network in the smaller box but a somewhat diffuse droplet in the larger box. Figure S9 shows that the mean junction separation of the networks for both simulation box sizes is independent of the network morphology and the presence of the periodic boundaries for the polymer lengths and concentrations studied.

### MOVIE LEGENDS

**Movie 1** Initial stage of the aggregation of 634 polymers $B_{16}$ (polymer concentration = 0.002) with low end-cap affinity $\varepsilon = 0.68$. This concentration is below the threshold for a stable network as shown quantitatively in Fig. 3 (blue curve, star symbols).

**Movie 2** Initial stage of the aggregation of 634 polymers $B_{16}$ with high end-cap affinity $\varepsilon = 0.8$. For this affinity, the network assembles rapidly as quantified in Fig. 3 (red curve, star symbols).

**Movie 3** Initial stage of the aggregation of 634 polymers $B_{16}$ with very high end-cap affinity $\varepsilon = 0.96$. For this affinity, the network assembles rapidly as quantified in Fig. 3 (green curve, star symbols).

**Movie 4** Initial stage of the aggregation of 1251 polymers $B_8$ with high affinity $\varepsilon = 0.8$. The polymers condense into a large network similar to the $B_{16}$ case.

**Movie 5** Initial stage of the aggregation of 1180 polymers $B_{24}$ with high affinity $\varepsilon = 0.8$. The polymers condense into a large network similar to the $B_{16}$ case.

### REFERENCES


1 R. D. Groot and P. B. Warren, "Dissipative Particle Dynamics: Bridging the Gap Between Atomistic and Mesoscopic Simulations," J. Chem. Phys. **107**, 4423-4435 (1997).
